# A community-curated, global atlas of *Bacillus cereus sensu lato* genomes for epidemiological surveillance

**DOI:** 10.1101/2023.12.20.572685

**Authors:** Vignesh Ramnath, Martin Larralde, Pedro Menchik, Ariel J. Buehler, Anna Sophia Harrand, Taejung Chung, Xiaoyuan Wei, Vishnu Raghuram, Hadrien Gourlé, Rian Pierneef, Itumeleng Matle, Marina Aspholm, Magnus Andersson, Rachel A. Cheng, Jasna Kovac, Johan Henriksson, Laura M. Carroll

## Abstract

The ability to cause foodborne illness, anthrax, and other infections has been attributed to numerous lineages within *Bacillus cereus sensu lato* (*s.l.*). However, existing pathogen surveillance databases facilitate dangerous pathogen misidentifications when applied to *B. cereus s.l.*, potentially hindering outbreak or bioterrorism attack response efforts. To address this, we developed BTyperDB (www.btyper.app), an atlas of *B. cereus s.l.* genomes with standardized, community-curated metadata. BTyperDB aggregates all publicly available *B. cereus s.l.* genomes (including >2,600 previously unassembled genomes) with novel genomes donated by laboratories around the world, nearly doubling the number of publicly available *B. cereus s.l.* genomes. To showcase its utility for pathogen surveillance, we use BTyperDB to identify emerging anthrax toxin- and capsule-harboring lineages. Overall, our study provides insight into the epidemiology of an under-studied group of emerging pathogens and highlights the benefits of inclusive, community-driven metadata FAIRification efforts.

---

*Bacillus cereus sensu lato* (*s.l.*), also known as the *B. cereus* group, is a complex of Gram-positive, environmentally widespread, spore forming species, which vary in terms of their industrial relevance and pathogenic potential.^1–3^ Some *B. cereus s.l.* strains play critical roles in agricultural or industrial settings (e.g., as environmentally friendly biopesticides, as prolific producers of antibiotics and other natural products, as food spoilage organisms that impact food security and sustainability).^3–8^ Other strains are pathogenic, and a range of illnesses have been attributed to *B. cereus s.l.*, including foodborne emetic intoxication, foodborne diarrheal toxicoinfection, severe non-gastrointestinal infections, and anthrax.^1–3,9^

Whole-genome sequencing (WGS) is being used increasingly by laboratories around the world to characterize *B. cereus s.l.* strains for a range of applications, including pathogen surveillance, food safety and spoilage risk evaluation, and novel biomolecule discovery.^3,6,10,11^ These genomes are deposited in public databases, such as the National Center for Biotechnology Information (NCBI) Assembly and Sequence Read Archive (SRA) databases, often with metadata related to the origin of the associated *B. cereus s.l.* strain. NCBI’s BioSample database, for example, links user-submitted metadata (e.g., taxonomic information, year or date of isolation, geographic location of isolation, host or isolation source) to WGS reads and/or assembled genomes deposited in the NCBI SRA and Assembly databases, respectively.^12–14^

In the pathogen surveillance realm, international WGS (meta)data sharing efforts have been shown to reduce the public health and economic burdens imposed by some pathogens.^15–18^ However, *B. cereus s.l.* poses several unique challenges in this space. First, because there is no single, universally accepted taxonomic framework for *B. cereus s.l.*, taxonomic metadata linked to publicly available *B. cereus s.l.* genomes are not standardized.^3^ Public databases are populated with *B. cereus s.l.* genomes with misleading, ambiguous, and/or incorrect species names, which can lead–and has led–to dangerous pathogen misidentifications and delayed response times in outbreak scenarios.^3,10,19^ Further, species names alone are inadequate for properly communicating *B. cereus s.l.* pathogenic potential, as several important *B. cereus s.l.* virulence factors are harbored on plasmids, which can be gained, lost, variably present within species, and present across multiple species (e.g., anthrax toxin genes *cya*, *lef*, and *pagA*; polyglutamate capsule genes *capBCADE*; cereulide [emetic toxin] synthetase-encoding *cesABCD*).^1,10,20,21^ A one-size-fits-all, species-centric approach to pathogen surveillance is thus inadequate for *B. cereus s.l.*, as strain-specific indicators of virulence potential go unaccounted for in existing databases (e.g., NCBI Pathogen Detection, https://www.ncbi.nlm.nih.gov/pathogens/).

Secondly, epidemiological surveillance of *B. cereus s.l.* is challenging, as illnesses caused by *B. cereus s.l.* are under-studied and underreported.^1,22^ *B. cereus s.l.* foodborne illness cases, for example, are typically self-limiting; thus, clinical samples are often not available, and when they are, the resulting *B. cereus s.l.* isolates are often not prioritized for WGS.^2,23^ Further, non-gastrointestinal infections caused by *B. cereus s.l.* are also likely under-sampled and under-sequenced, as *B. cereus s.l.* is often viewed as an environmental or laboratory contaminant, rather than a potential cause of these infections.^2^ Even when WGS data is deposited publicly, it is often in the form of sequencing reads that are never assembled into genomes. As a result, these unassembled *B. cereus s.l.* genomes are currently not included in WGS-based pathogen surveillance efforts.^10,14,24–26^

Finally, user-submitted metadata associated with publicly available pathogen genomes (including those of *B. cereus s.l.*) are often incomplete, unstandardized, and/or of poor quality. This makes it challenging, time-consuming, and sometimes impossible for WGS (meta)data to be reused and repurposed for epidemiological investigations.^27–29^ For example, in NCBI Pathogen Detection, over 25% of foodborne pathogen genomes lack key metadata attributes.^27^ Alarmingly, many historically important *B. cereus s.l.* strains (e.g., those linked to outbreaks, illness cases, bioterrorism attacks) do not have metadata available in their BioSample records (Supplementary Fig. S1). As a result, pathogen surveillance efforts that rely solely on BioSample entries for metadata (e.g., NCBI Pathogen Detection) are omitting critical information that can be used in *B. cereus s.l.* surveillance.^12^

Taken together, (i) taxonomic ambiguity, (ii) epidemiological and microbiological knowledge/data gaps, and (iii) incomplete metadata have the potential to hinder public health response efforts in *B. cereus s.l.* outbreak or bioterrorism attack scenarios.^10,19,27^ To address these critical gaps, we developed BTyperDB (www.btyper.app), an atlas of *B. cereus s.l.* genomes with standardized, community-curated metadata (Fig. 1). By aggregating all publicly available *B. cereus s.l.* genomes (including >2,600 previously unassembled genomes) with novel genomes generously donated by laboratories around the world, BTyperDB nearly doubles the number of *B. cereus s.l.* genomes available for pathogen surveillance and source tracking. Further, extensive metadata aggregation and curation efforts undertaken by *B. cereus s.l.* researchers and stakeholders around the globe have allowed us to populate BTyperDB with standardized metadata that can be leveraged in epidemiological investigations. Overall, BTyperDB represents a community-curated, genomic atlas, which can be used to provide much-needed insight into the epidemiology and biology of emerging pathogens within *B. cereus s.l*.

**Fig. 1.**
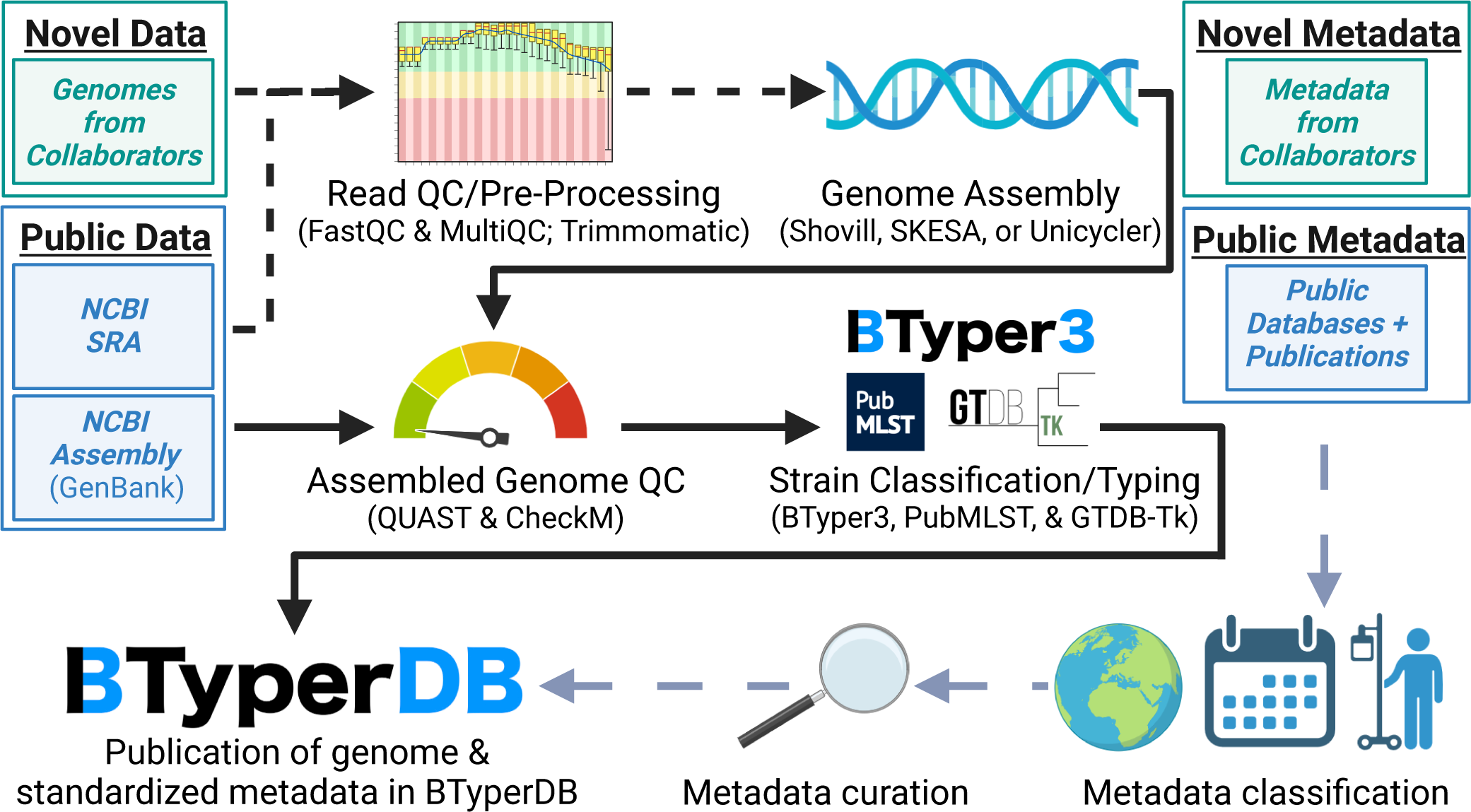
Overview of the BTyperDB workflow. Briefly, genomes represented by sequencing reads (previously unassembled genomes from NCBI’s Sequence Read Archive [SRA] database, or genomes donated by collaborators) undergo quality control (QC) and pre-processing steps and are assembled into contigs. Assembled genomes (those generated from sequencing reads in this study, plus previously assembled GenBank genomes from NCBI’s Assembly database) undergo QC and taxonomic assignment/sequence typing using a variety of methods. Genomes of adequate quality are deposited in BTyperDB alongside metadata (provided by collaborators or obtained via a search of public databases and/or publications), which has been manually curated and standardized using the BTyperDB Ontology (a *B. cereus s.l.*-specific ontology).

## RESULTS

### BTyperDB nearly doubles the number of *B. cereus s.l.* genomes available for pathogen surveillance

To construct BTyperDB, we first downloaded all publicly available, assembled *B. cereus s.l.* genomes from NCBI’s Assembly database (*n* = 3,428 GenBank assemblies; Fig. 1). Because *B. cereus s.l.* WGS data is often deposited in public databases as unassembled sequencing reads, we further downloaded all previously unassembled *B. cereus s.l.* genomes in NCBI’s SRA database and constructed assemblies for each (*n* = 3,398 previously unassembled genomes from NCBI SRA; Fig. 1). Finally, several laboratories around the world generously donated novel *B. cereus s.l.* genomes, which had previously never been publicly available (*n* = 75 previously unpublished genomes; Fig. 1). Together, after filtering out low-quality genomes, the WGS data generation, assembly, and aggregation efforts undertaken here allowed us to nearly double the number of publicly available, assembled *B. cereus s.l.* genomes, while more than quadrupling and tripling the number of anthrax toxin-harboring and cereulide (emetic toxin) synthetase-harboring genomes, respectively (see the “Methods” section below for details; Fig. 1 and 2a, Supplementary Table S1).

**Fig. 2.**
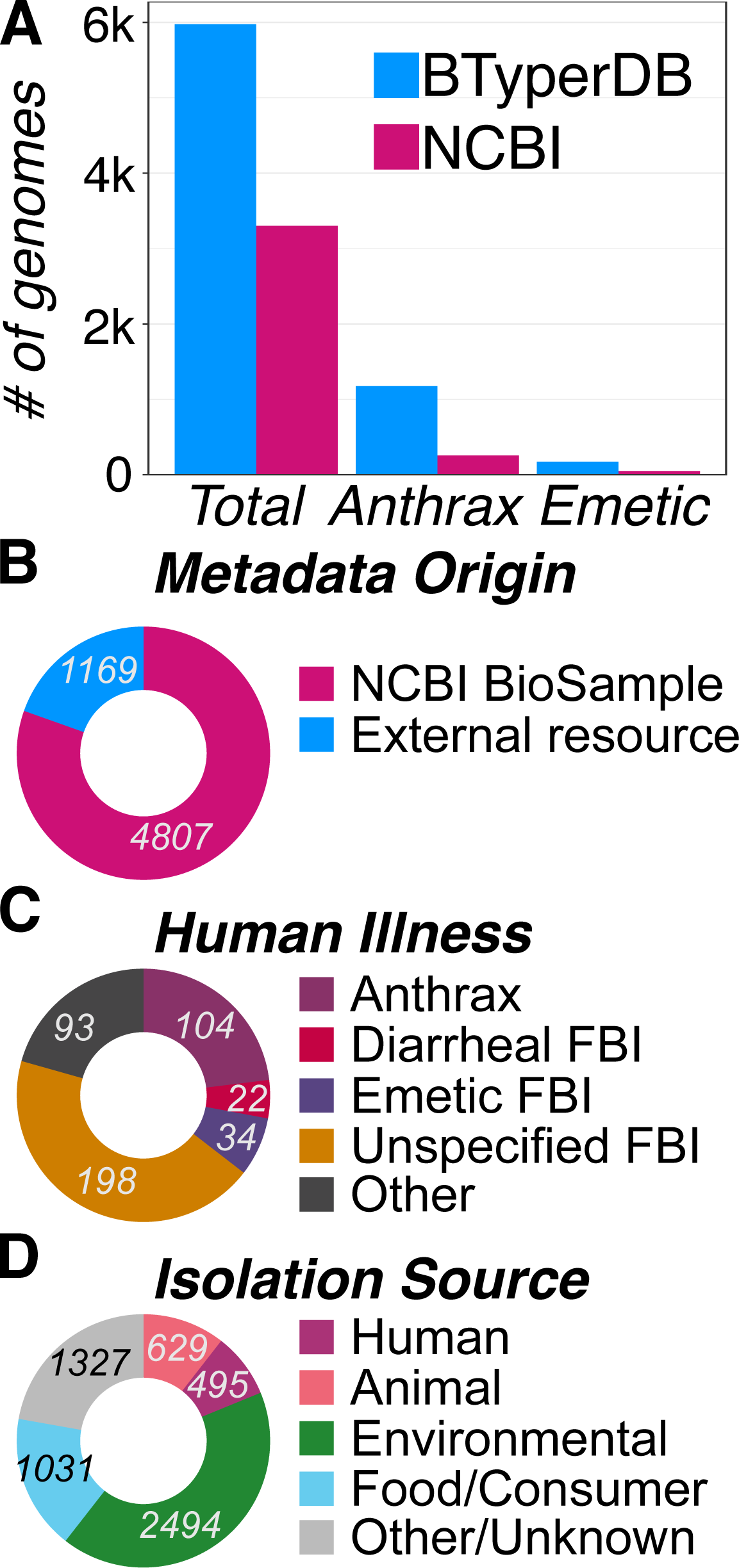
Overview of all *B. cereus s.l.* genomes in BTyperDB and their corresponding curated, standardized metadata. (A) Total number of assembled *B. cereus s.l.* genomes in NCBI (pink) and BTyperDB (blue; “Total”), including genomes with (i) anthrax toxin genes (≥1 of *cya*, *lef*, and/or *pagA*; “Anthrax”) or (ii) cereulide (emetic toxin) synthetase genes (all of *cesABCD*; “Emetic”). (B) Origins of metadata in BTyperDB. For 80% of genomes, all metadata attributes were retrieved from NCBI’s BioSample database (4,807 of 5,976 genomes, 80.4%; pink). However, for 20% of genomes, external resources (e.g., peer-reviewed publications) provided additional metadata, which was absent from NCBI but included in BTyperDB (1,169 of 5,976 genomes, 19.6%; blue). Proportion of BTyperDB genomes isolated (C) in conjunction with a human illness case, or (D) from a given source (the BTyperDB Ontology “Human_Illness” and “Source_1” attributes, respectively). For B, C, and D, numeric labels denote the number of genomes assigned to a given metadata source, illness, or isolation source, respectively. For (C), genomes categorized as “Unknown” are excluded. FBI, foodborne illness; Consumer, consumer product.

Notably, there is no single, universally accepted taxonomy for *B. cereus s.l.^3^* Thus, we subjected each genome in BTyperDB to multiple species assignment and sequence typing methods, so that users can search for and download genomes using standardized taxonomic labels assigned via their preferred method(s) (Fig. 1, Supplementary Table S1). Further, genomospecies assignments are inadequate for conveying *B. cereus s.l.* pathogenic potential at the strain level, as many important *B. cereus s.l.* virulence factors can be gained, lost, and are variably present within and across species boundaries.^10^ To overcome this, BTyperDB allows users to query genomes based on the presence and absence of *B. cereus s.l.* virulence factors, including genes encoding anthrax toxin, capsules, cereulide (emetic toxin) synthetase, and diarrheal enterotoxins (see “Taxonomic assignment and sequence typing” below for details; Fig. 1 and 2a, Supplementary Table S1).

Finally, alarmingly little is known about the epidemiology of illnesses caused by *B. cereus s.l.* on a large scale (foodborne illnesses, in particular).^3,30^ To date, there has been no way to systematically identify which *B. cereus s.l.* genomes are derived from strains linked to human illness cases or outbreaks, let alone strains linked to specific types of illness (e.g., diarrheal toxicoinfection). To that end, we manually searched for clinical and epidemiological data linked to each genome in BTyperDB, first by querying each genome’s BioSample record, and then by reviewing external sources (e.g., peer-reviewed publications; Fig. 2b). Metadata for each of the 5,976 high-quality genomes was standardized using the BTyperDB Ontology, a *B. cereus s.l.*-specific ontology, which we developed (Fig. 2cd, Supplementary Table S1). As a result, users can rapidly identify *B. cereus s.l.* strains isolated from various sources (e.g., human-, animal-, food-associated; Fig. 2d) and geographic regions (e.g., continent, country, first-level administrative division; Fig. 3, Supplementary Fig. S2-S6). Further, for strains linked to human clinical cases, users can search by clinical manifestation (e.g., anthrax, diarrheal illness; Fig 2c), as well as identify groups of strains linked to specific human outbreaks using unique outbreak identifiers (Fig. 4, Supplementary Fig. S2-S6, Supplementary Table S1).

**Fig. 3.**
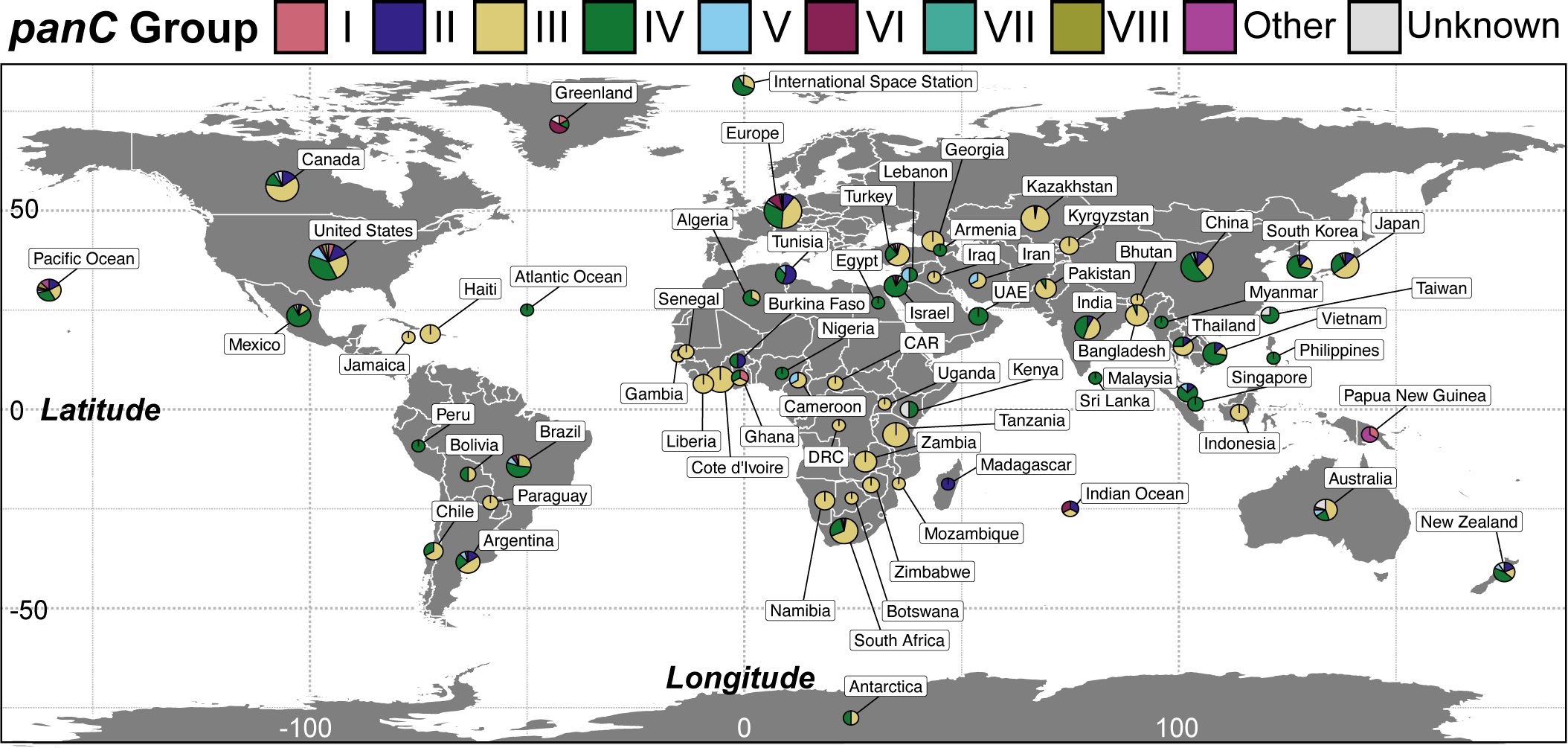
Country of origin for all *B. cereus s.l.* genomes in BTyperDB. Pie charts are sized according to the number of genomes from a particular country and show the proportion of genomes assigned to each *panC* phylogenetic group (i.e., historically important “species”-like units delineated using the sequence of *panC*). For readability, all genomes from “Europe” (as defined via pycountry) are aggregated into a single pie chart. For related figures, see Supplementary Fig. S3-S5.

**Fig. 4.**
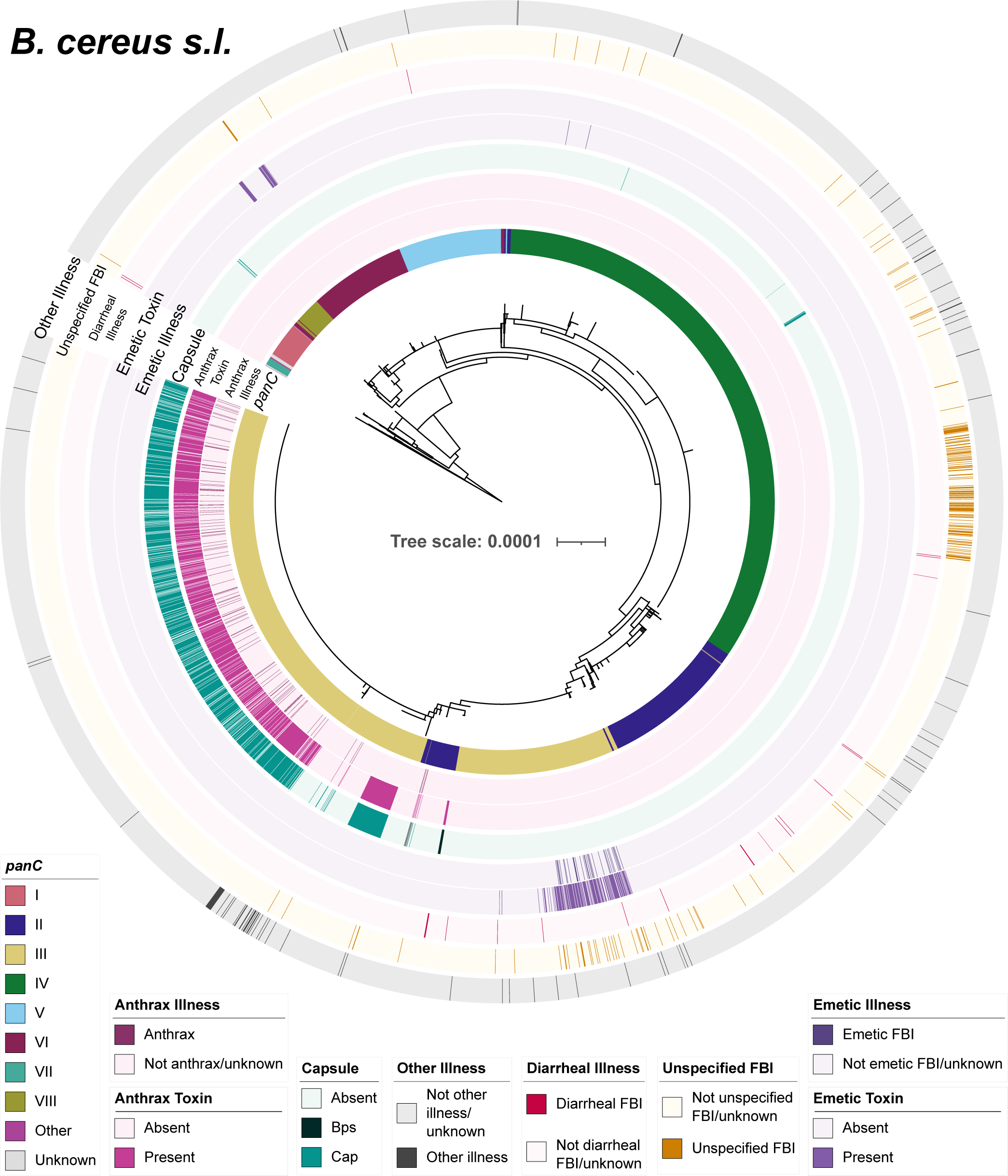
Clinical metadata and selected virulence factors present in 5,976 BTyperDB genomes. Color strips surrounding the maximum likelihood phylogeny denote (from interior to exterior): (i) *panC* phylogenetic group (i.e., historically important “species”-like units delineated using the sequence of *panC*); (ii) whether a genome was linked to a human anthrax case or not; (iii) presence and absence of two or more anthrax toxin genes (*cya*, *lef*, and/or *pagA*); (iv) presence and absence of select capsular genes (all of *capBCADE* or *bpsXABCDEFGH*); (v) whether a genome was linked to a human foodborne emetic intoxication case or not; (vi) presence and absence of emetic toxin (cereulide) synthetase-encoding *cesABCD*; whether a genome was linked to a (vii) human diarrheal illness case, (viii) unspecified human foodborne illness (FBI) case, or (ix) other human illness/infection, or not. The phylogeny is rooted using an outgroup (omitted for readability), with branch lengths in substitutions per site. For an extended version of this phylogeny, see Supplementary Fig. S6.

Altogether, the extensive community-driven WGS (meta)data acquisition and curation efforts conducted here (i) nearly double the number of publicly available *B. cereus s.l.* genomes, and (ii) provide standardized metadata that link *B. cereus s.l.* genomes to their respective clinical and epidemiological manifestations on an unprecedented scale (Fig. 1-4, Supplementary Fig. S2-S6). To showcase its utility, we use BTyperDB to identify and track the global spread of novel, emerging *B. cereus s.l.* lineages, which have gained the ability to produce anthrax-associated virulence factors, including (i) a novel anthrax toxin gene-harboring lineage responsible for diarrheal illness in China, and (ii) polyglutamate capsule-encoding lineages isolated from insect food products and human clinical cases. Each case study is described in detail below.

### A novel anthrax toxin gene-harboring *B. tropicus* lineage responsible for diarrheal illness in China harbors rare, exo-polysaccharide-encoding genes

Predicting whether a *B. cereus s.l.* strain is capable of causing diarrheal illness or not is a major open problem in the food safety space,^3,31,32^ due in part to the fact that nearly all *B. cereus s.l.* strains possess enterotoxin genes,^20^ a result confirmed here (Supplementary Table S1). This, combined with poor epidemiological surveillance of the disease, means that it is unclear which *B. cereus s.l.* strains are “high-risk” in terms of their ability to cause diarrheal illness.

To address this critical knowledge gap in *B. cereus s.l.* epidemiology, we manually searched for diarrheal illness cases linked to nearly 6,000 genomes in BTyperDB. Overall, we identified 22 *B. cereus s.l.* genomes reportedly isolated in conjunction with diarrheal illness cases in humans (Fig. 2b, 4; Supplementary Fig. S4-S6, Supplementary Table S1). Notably, the 22 genomes spanned multiple *B. cereus s.l.* species, regardless of which taxonomic framework was used (Fig. 4, Supplementary Fig. S6, Supplementary Table S1). Further, the phylogenetic groups (i.e., historically important “species”-like units delineated using the sequence of *panC*, referred to hereafter as “*panC* Groups”)^33^ attributed to the most diarrheal foodborne illness cases were *panC* Groups II and III, together accounting for nearly three-fourths of all *B. cereus s.l.* genomes linked to diarrheal illness cases (*n* = 11 and 5 genomes, 50% and 23% of 22 genomes, respectively; Fig. 4, Supplementary Fig. S6, Supplementary Table S1).

Interestingly, three *panC* Group II strains assembled in this study, which were reportedly responsible for diarrheal illness in 2017 in Jiangxi, China, harbored anthrax toxin genes *cya* and *pagA*, which encode the anthrax edema factor (EF) and protective antigen (PA), respectively (*lef*, which encodes the anthrax lethal factor [LF], was not detected; Fig. 4, 5). In addition, all three genomes possessed *bpsXABCDEFGH*, which together encode a rare, antiphagocytic exopolysaccharide capsule that enables some anthrax-causing strains with “*B. cereus*”-like phenotypic characteristics (per the U.S. Food and Drug Administration Bacteriological Analytical Manual; FDA BAM)^34^ to evade the host immune system (Fig. 4, 5, Supplementary Table S1). Notably, all three genomes belonged to sequence type 234 (ST234), a ST within the *B. tropicus* genomospecies (defined via the Genome Taxonomy Database [GTDB]; Fig. 5, Supplementary Fig. S7, S8).

**Fig. 5.**
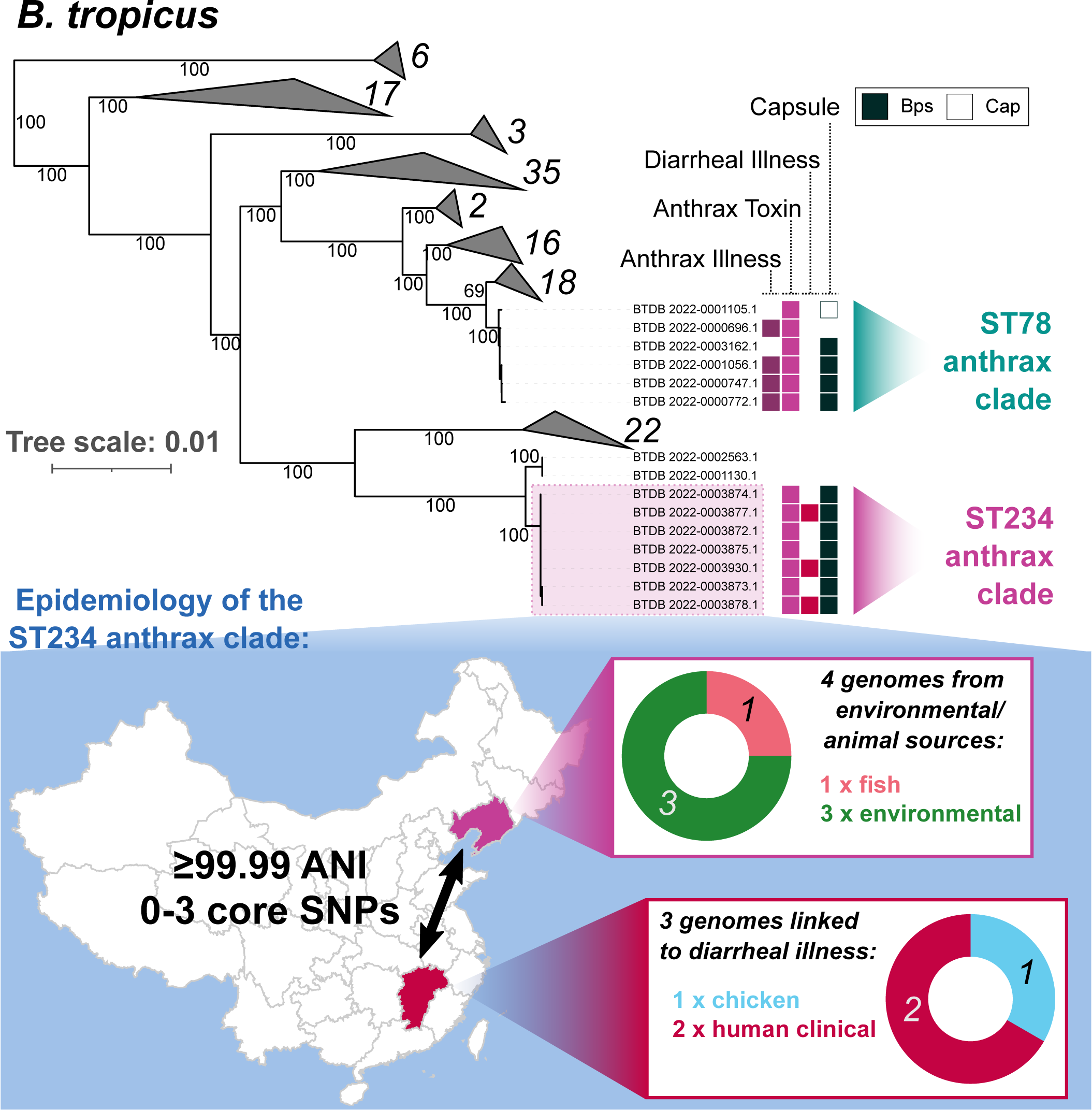
A novel anthrax toxin gene-harboring *B. tropicus* lineage responsible for diarrheal illness in China harbors genes encoding the rare Bps exo-polysaccharide capsule (highlighted in the map and labeled in the maximum likelihood phylogeny as “ST234 anthrax clade”; *n* = 7 genomes, ultrafast bootstrap = 100%). The seven genomes within the ST234 anthrax clade were isolated from Liaoning Province and Jiangxi Province (four and three genomes, respectively) in 2017 and were nearly identical (≥99.99 ANI, difference of 0-3 core SNPs, presence/absence of 0-5 accessory genes; see the map below the phylogeny). The three genomes linked to diarrheal illness harbored anthrax toxin genes *cya* and *pagA*; *lef* could not be detected in any of the three genomes at >50% amino acid identity and >40% coverage. The maximum likelihood phylogeny was constructed using all genomes assigned to *B. tropicus* (per GTDB; *n* = 134). Colored shapes to the right of the phylogeny (from left to right) denote genomes: (i) linked to a human anthrax case; (ii) with two or more anthrax toxin genes (*cya*, *lef*, and/or *pagA*); (iii) linked to a human diarrheal illness case; (iv) with select capsular genes (all of *capBCADE* or *bpsXABCDEFGH*; Cap and Bps, respectively). Clades that did not contain anthrax toxin or capsular genes are collapsed, with numerical labels denoting the number of genomes in each. Branch labels denote branch support percentages obtained using 1,000 replicates of the ultrafast bootstrap approximation. The phylogeny is rooted using an outgroup (omitted for readability), with branch lengths in substitutions per site. Donut charts below the phylogeny showcase the proportion of genomes from each highlighted province assigned to each source; numeric labels within the donut chart denote the number of genomes assigned to that source. For related figures, see Supplementary Fig. S7-S10.

While anthrax-causing *B. tropicus* genomes have been described before,^26^ they are incredibly rare. In a 2022 study querying all assembled *B. cereus s.l.* genomes in NCBI,^26^ anthrax toxin genes were confined to ST78 strains from two geographic regions: (i) the U.S. Gulf Coast region, from which four anthrax-causing human clinical strains originated, and (ii) China, which was home to a single genome (isolated from a kangaroo; Fig. 5, Supplementary Fig. S7-S9).^26^ Importantly, in the 2022 study,^26^ *bps* exo-polysaccharide capsule-encoding genes were confined to the U.S. Gulf Coast; thus, the three ST234 genomes assembled and identified here not only represent a novel anthrax toxin gene-harboring lineage within *B. tropicus*, but also indicate that the rare, antiphagocytic *bps* exo-polysaccharide capsule exhibits a wider geographic spread than previously thought (Fig. 5, Supplementary Fig. S7-S10).

To gain insight into the emergence and subsequent dissemination of rare, anthrax toxin gene-harboring *B. tropicus* lineages, we used BTyperDB to download all *B. tropicus* genomes (per GTDB’s taxonomy), including all members of ST234 (Fig. 5, Supplementary Fig. S7-S10, Supplementary Table S1). Notably, all anthrax toxin gene-harboring ST234 genomes (including the three reportedly linked to diarrheal illness) were isolated in 2017 in China, possessed *bps* exo-polysaccharide capsule-encoding genes, and were extremely closely related to each other, indicating that the presence of anthrax toxin and *bps* genes in this lineage was the result of one or more acquisition event(s) (Fig. 5, Supplementary Fig. S10, Supplementary Table S1).

Interestingly, the three genomes responsible for diarrheal illness were reportedly isolated from human clinical samples and food (chicken) in Jiangxi Province; however, it is unclear from the metadata whether the reported diarrheal illnesses were caused by enterotoxins or anthrax toxin (Fig. 5). Four additional anthrax toxin gene-harboring ST234 genomes were isolated from groundwater, wastewater, fish, and a probiotic product in Liaoning Province, indicating that members of this closely related, anthrax toxin- and *bps* -harboring lineage are distributed across multiple Chinese provinces (Fig. 5, Supplementary Fig. S10, Supplementary Table S1).

Overall, our findings highlight the importance of moving beyond species names when evaluating *B. cereus s.l.* public health and food safety risks. Multiple *B. cereus s.l.* species have been isolated in conjunction with diarrheal illness and anthrax cases, and strains within and across these species showcase heterogeneity in terms of the virulence factors they possess. Therefore, *B. cereus s.l.* species should not be treated as single, homogenous units for risk evaluation purposes, particularly when *B. cereus s.l.* taxonomy is so hotly contested.

### Polyglutamate capsule-encoding genes can be variably acquired within and across *B. cereus s.l.* species boundaries

In addition to the anthrax toxins (i.e., PA, LF, EF), the antiphagocytic polyglutamate capsule produced by *B. anthracis^34^* is considered one of two principal anthrax virulence factors, as it equips producer strains with the ability to evade the host immune system during anthrax infection.^35,36^ While historically viewed as a species-defining characteristic of “*B. anthracis*”, polyglutamate capsule-encoding *capBCADE* are typically located on a plasmid and are variably present within the *B. anthracis* genomospecies.^26^ On rare occasions, *capBCADE*-harboring strains outside of *B. anthracis* have been described.^26,37,38^ However, the epidemiology of *capBCADE*-harboring *B. cereus s.l.* has not been evaluated systematically, and the clinical relevance of these strains is not understood (particularly when anthrax toxin genes are absent).

To gain insight into the global epidemiology and virulence potential of *capBCADE*-harboring *B. cereus s.l.*, we queried BTyperDB for all *capBCADE*-harboring genomes (Fig. 4, Supplementary Table S1). Notably, while largely confined to the *B. anthracis* genomospecies (per GTDB’s definition, 1,133 of 1,148 genomes, 98.7%), *capBCADE* were more taxonomically widespread than previously thought: aside from one anthrax toxin-gene harboring *B. tropicus* genome (i.e., the strain isolated from a kangaroo in China mentioned above; Fig. 5), *capBCADE* were identified in *B. cereus*, *B. nitratireducens*, and *B. thuringiensis* (per GTDB’s definitions, 9, 4, and 1 genomes, respectively; Fig. 4, Supplementary Table S1). Importantly, none of these 14 *capBCADE*-harboring genomes possessed anthrax toxin encoding-genes and thus do not represent an anthrax threat (Fig. 4, Supplementary Table S1).

Interestingly, all nine *panC* Group IV *capBCADE*-harboring genomes (i.e., all *capBCADE*-harboring genomes classified as “*B. cereus*”, plus the lone *capBCADE*-harboring “*B. thuringiensis*” genome, per GTDB), were reportedly isolated from insect-based food products in Germany in 2020 (Fig. 6a, Supplementary Fig. S11, Supplementary Table S1). Despite this, the *capBCADE*-harboring *panC* group IV genomes were considerably diverse, indicating that the presence of polyglutamate capsule-encoding genes within this species unit is the result of multiple separate acquisition events (Fig. 6a, Supplementary Fig. S11).

**Fig. 6.**
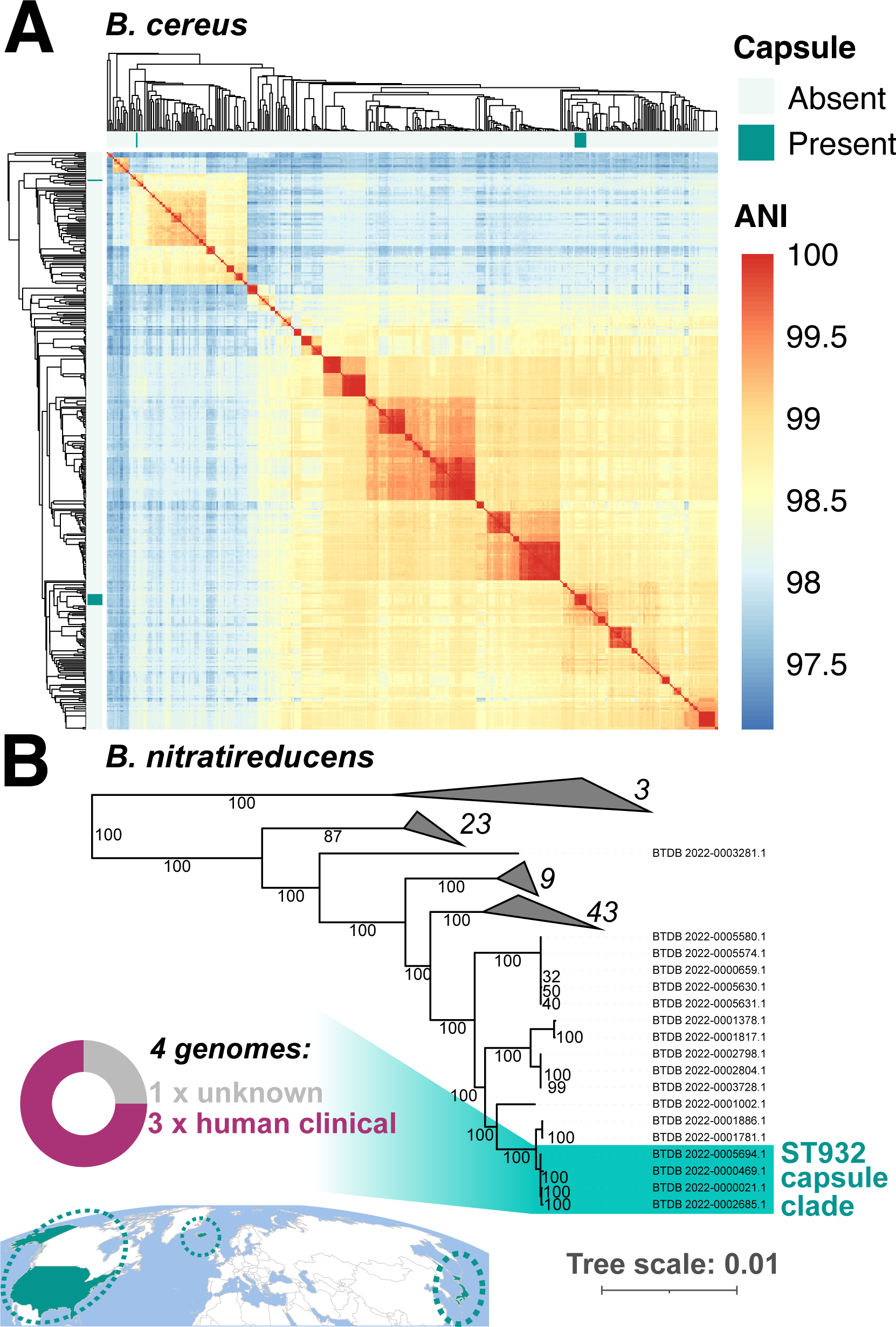
Polyglutamate capsule genes are variably present within and across non-*anthracis B. cereus s.l.* species. (A) Heatmap of pairwise average nucleotide identity (ANI) values calculated between all *B. cereus* genomes (per GTDB; *n* = 418 genomes). Presence and absence of polyglutamate capsule-encoding *capBCADE* in each genome are displayed in the color strips surrounding the heatmap. Dendrograms were produced via average linkage hierarchical clustering of genomes using ANI dissimilarities (100-ANI). Notably, multiple *B. cereus* lineages from insect-based food products have acquired polyglutamate capsule-encoding genes: eight of nine genomes shared ≥99.9 ANI, differed by 8-136 core SNPs, and formed a well-supported clade within the *B. cereus* phylogeny (ultrafast bootstrap = 100%, Supplementary Fig. S11). (B) Polyglutamate capsule-harboring *B. nitratireducens* are confined to the ST932 lineage (highlighted and labeled in the maximum likelihood phylogeny as “ST932 capsule clade”; *n* = 4 genomes, ultrafast bootstrap = 100%). Within the ST932 capsule clade, two genomes from Japan were nearly identical (>99.9 ANI, 0 core SNPs difference), but the clade as a whole was more diverse (99.6-100.0 ANI, difference of 0-311 core SNPs). The phylogeny was constructed using all *B. nitratireducens* genomes (per GTDB; *n* = 96). The map below the phylogeny denotes countries in which members of the ST932 capsule clade have been detected (i.e., Japan, Iceland, and the United States); the donut chart contains reported isolation sources for each ST932 capsule clade genome. Branch labels denote branch support percentages obtained using 1,000 replicates of the ultrafast bootstrap approximation. The phylogeny is rooted using an outgroup (omitted for readability), with branch lengths in substitutions per site. Selected clades are collapsed for readability, with numerical labels denoting the number of genomes in each. For related figures, see Supplementary Fig. S12-S14.

Comparatively, *capBCADE*-harboring *B. nitratireducens* all belonged to a single lineage, ST932 (Fig. 6b, Supplementary Fig. S12-S14, Supplementary Table S1). Within ST932, all four *capBCADE*-harboring strains formed a well-supported clade (Fig. 6b, Supplementary Fig. S13, S14). Two genomes from Japan were nearly identical, but the clade as a whole was more diverse and spanned three countries on three continents (Fig. 6b, Supplementary Fig. S12-S14, Supplementary Table S1). Interestingly, aside from one Japanese strain of unknown origin, the three remaining *capBCADE*-harboring *B. nitratireducens* were isolated from human clinical cases (one from each of Japan, Iceland, and the United States), although it is unclear what role the polyglutamate capsule played in these infections, if any (Fig. 6b, Supplementary Fig. S12-S14).

Altogether, these results highlight the importance of accounting for plasmid-mediated virulence factors in *B. cereus s.l.* genomic surveillance. As demonstrated above, *B. cereus s.l.* lineages (i.e., distinct phylogenetic clades within species, as well as individual strains within a clade) can acquire novel virulence characteristics, which allow them to cause illness and/or interact with the human host. Thus, the strain-specific surveillance approaches implemented in BTyperDB can be used to identify the emergence and monitor the spread of novel, high-risk lineages within *B. cereus s.l*.

## DISCUSSION

Public databases, data hubs, and data portals that link pathogen WGS data to clinically and epidemiologically relevant metadata have become essential resources for pathogen surveillance in the genomic era.^12,27,39^ The WGS (meta)data sharing efforts facilitated by these resources have been shown to reduce the time it takes to solve and respond to outbreaks, leading to better public health outcomes and lower burdens of illness for priority pathogens.^15,16,27,40^ Historically, *B. cereus s.l.* has been overlooked by genomic surveillance efforts, despite the fact that the species complex contains bioterrorism agents, foodborne pathogens, and industrially relevant microbes; even when *B. cereus s.l.* is considered, existing tools (e.g., NCBI Pathogen Detection) are inadequate for *B. cereus s.l.* surveillance, as they do not account for within- and between-species heterogeneity in virulence potential, nor do they provide taxonomic assignments using standardized, reproducible methods.

Simultaneously, because it is under-sampled and under-sequenced, *B. cereus s.l.* represents an unprecedented opportunity to showcase the value of community-driven (i) WGS data generation and (ii) metadata curation and standardization efforts. Specifically, calls to “FAIRify” pathogen (meta)data (i.e., making [meta]data “Findable”, “Accessible”, “Interoperable”, and “Reusable”) have been made, but this task remains daunting, due to the sheer number of genomes available for priority pathogens.^12,27^ Additionally, important domain-specific metadata (e.g., pathogen-specific attributes, such as the ability to cause a particular illness, the ability to produce a specific toxin, pathogen-specific phenotypes) are often omitted from general-purpose pathogen surveillance databases, as such information is often difficult to standardize.^28^ To construct BTyperDB, *B. cereus s.l.* researchers, public health officials, and industry professionals around the world collectively spent thousands of hours aggregating, curating, and standardizing metadata for every *B. cereus s.l.* genome available at the time of the data freeze–including metadata from “unfindable” sources (e.g., supplementary materials of published manuscripts, historical manuscripts). Further, the extensive genome assembly efforts undertaken here, including the assembly of dozens of previously unpublished genomes generously donated by laboratories around the world, nearly double the number of publicly available *B. cereus s.l.* genomes, which can be used in pathogen surveillance efforts.

Despite the massive (meta)data curation and standardization efforts undertaken here, it is imperative to point out the limitations of BTyperDB and public pathogen genome collections more broadly. First and foremost, public pathogen genome collections are at the mercy of user-submitted metadata. Any mistakes that users make when uploading genomes or metadata can get propagated to downstream databases. Here, we not only utilized metadata in NCBI’s BioSample database, but we additionally conducted a literature search for every *B. cereus s.l.* genome available in NCBI, with potential data provenance issues noted by curators (Supplementary Table S1). However, even with these extra precautions, user mistakes in metadata reporting can still make their way into databases (e.g., due to mistakes in publications). Second, some metadata attributes are often unavailable in public databases and not reported in publications. For example, *B. cereus s.l.* strains from human clinical cases are reported as such, but the specific type of illness (e.g., emetic intoxication, diarrheal toxicoinfection) is often not specified. This is often not the user’s fault; as mentioned above, pathogen-specific metadata are often not captured in general-purpose databases,^28^ and we anticipate that domain-specific surveillance tools like BTyperDB can help remedy this. Finally, just because a pathogen was isolated in conjunction with an illness or outbreak does not mean that particular strain was responsible for the illness case(s). These scenarios have been observed for other pathogens (e.g., *Staphylococcus aureus*),^41^ but they are particularly relevant for *B. cereus s.l.*, as environmental strains can be isolated from human clinical samples and foods and incorrectly classified as “human clinical” strains by users.^23^ Additional epidemiological, microbiological, and clinical metadata are needed to link *B. cereus s.l.* genomes to human illness cases, and we anticipate that BTyperDB represents an important first step toward enabling the collection and reporting of such metadata.

Altogether, BTyperDB is the culmination of a massive (meta)data FAIRification effort undertaken by the community generating and utilizing that (meta)data. As a resource, BTyperDB will allow scientists and other stakeholders (including those with limited bioinformatics experience) to easily acquire massive amounts of relevant WGS data, enabling them to (i) rapidly investigate outbreaks and illness cases, (ii) track pathogen evolution and spread, (iii) identify genomic determinants associated with phenotypes of interest (e.g., via genome-wide association studies), and (iv) identify novel, emerging lineages. Our efforts further highlight the importance of incorporating domain-specific knowledge into pathogen surveillance settings and provide a flexible framework to enable future *B. cereus s.l.* metadata collection and reporting efforts. Finally, the efforts undertaken here showcase the commitment of the *B. cereus s.l.* community to FAIR principles in genomic surveillance settings.

## METHODS

### Acquisition and assembly of putative *B. cereus s.l.* genomes from publicly available genomic data

All child nodes of NCBI’s “*Bacillus cereus* group” taxonomic level (NCBI Taxonomy ID 86661) were treated as potential *B. cereus s.l.* members, and a list of their NCBI Taxonomy IDs was compiled (https://www.ncbi.nlm.nih.gov/taxonomy/?term=txid86661[Subtree], accessed 8 August 2022).^42^ To ensure that no validly published and effective *B. cereus s.l.* species were absent from the list of *B. cereus s.l.* Taxonomy IDs prepared by NCBI, all validly published and effective *B. cereus s.l.* species names were queried and manually added if not originally present in NCBI’s *B. cereus s.l.* list (*n* = 28 species, accessed 8 August 2022; Supplementary Table S2, Supplementary Text).

All assembled GenBank genomes available in NCBI’s Assembly database (accessed 8 August 2022) were treated as candidate *B. cereus s.l.* genomes if the associated NCBI Taxonomy ID listed in the GenBank assembly summary file (https://ftp.ncbi.nlm.nih.gov/genomes/ASSEMBLY_REPORTS/assembly_summary_gen <u>bank.txt</u>; accessed 8 August 2022) was included in the *B. cereus s.l.* Taxonomy ID list compiled here (Supplementary Table S2). All candidate *B. cereus s.l.* assembled genomes were downloaded from GenBank via NCBI’s File Transfer Protocol (FTP) site (https://ftp.ncbi.nlm.nih.gov/; accessed 8 August 2022).

To search NCBI’s Sequence Read Archive (SRA) database^14^ for sequencing reads derived from potential *B. cereus s.l.* members, metadata in SRA was queried using Google Cloud’s gcloud command line interface (CLI, accessed 8 August 2022; Supplementary Text). SRA accession numbers were considered to belong to potential *B. cereus s.l.* members if the value in SRA’s “organism” field was equivalent to an organism name associated with one of the NCBI Taxonomy IDs compiled here (Supplementary Table S2, Supplementary Text).

Using the compiled list of potential *B. cereus s.l.* genomes available in NCBI’s (i) Assembly (GenBank) and (ii) SRA databases, a final list was compiled by taking the union of the NCBI BioSample^12^ accession numbers associated with each genome or SRA run, respectively (accessed 8 August 2022). For BioSamples with an associated assembled genome, the assembled GenBank genome available in the Assembly database was used in subsequent steps. For BioSamples with no associated assembled genome (i.e., those with SRA data alone), FASTQ files associated with the BioSample were downloaded using the SRA Toolkit v2.11.0 “prefetch” and “fastq-dump” commands.^14^ The quality of the resulting raw FASTQ files was assessed using FastQC v0.11.9^43^ and MultiQC v1.12.^44^ For SRA runs associated with paired-end Illumina reads, Shovill v1.1.0 (https://github.com/tseemann/shovill) was used to assemble each set of paired-end reads into contigs (Supplementary Text). For SRA runs linked to single-end Illumina reads, single-end reads were trimmed and assembled into contigs using Shovill’s implementation of Trimmomatic^45^ and SKESA,^46^ respectively (Supplementary Text). For SRA runs with long reads available (i.e., PacBio or MinION), Unicycler v0.5.0^47^ was used to assemble the genome; when paired-end Illumina reads were additionally available, both long and short reads were supplied as input to Unicycler, and a hybrid assembly was constructed (Supplementary Text). For SRA runs with paired-end Illumina and Roche 454 reads available, the paired-end Illumina reads were assembled into contigs using Shovill (Supplementary Text), with 454 reads omitted.

Overall, the genome aggregation and assembly efforts conducted here resulted in 6,826 candidate *B. cereus s.l.* genomes, which were used in subsequent steps: (i) 3,428 previously assembled, publicly available genomes from NCBI’s GenBank Assembly database, and (ii) 3,398 genomes assembled here from publicly available sequencing reads.

Whole-genome sequencing and assembly of South African *B. cereus s.l.* genomes.

The genomes of two *B. cereus s.l.* strains isolated from beef products in South Africa were additionally sequenced in this study (Supplementary Table S1). Briefly, the *B. cereus s.l.* strains were isolated from meat and meat products following the ISO 7932:2004 protocol, as described previously.^48^ Meat samples were homogenized in maximum recovery diluent (1:10 w/v), with subsequent 10-fold dilutions up to 10^−5^. Each dilution was surface-inoculated onto Mannitol Yolk Polymyxin (MYP) agar in duplicate plates and then incubated at 30 ± 1°C for 18−24 h. If bacterial colonies were not clearly visible, the MYP plates were further incubated for an additional 24 h. Presumptive *B. cereus s.l.* colonies, characterized by mannitol-negative appearance (pink to red) with a precipitate zone around the colonies (indicating lecithinase-positive reaction), were identified. These presumptive colonies were streaked onto Sheep Blood Agar to assess hemolysis, and those exhibiting beta-hemolysis were considered presumptive *B. cereus s.l*. Further confirmation involved subjecting presumptive *B. cereus s.l.* colonies to various biochemical tests, including acid production from phenol red glucose broth, nitrate reduction, acetylmethyl-carbinol production using Voges Proskauer (VP) medium, and acid production from mannitol.

The extraction of genomic DNA was performed using the High Pure PCR template preparation kit (Roche, Germany) as described previously.^48^ WGS of the isolates was carried out at the Biotechnology Platform, Agricultural Research Council, Onderstepoort, South Africa. DNA libraries were constructed using TruSeq and Nextera DNA library preparation kits (Illumina, San Diego, CA, USA), and subsequently subjected to sequencing on HiSeq and MiSeq instruments (Illumina, San Diego, CA, USA). The resulting paired-end Illumina reads associated with each strain were trimmed via Trimmomatic v0.38 (Supplementary Text). The quality of the resulting trimmed paired-end reads was assessed using FastQC, and Shovill was used to assemble each genome (Supplementary Text). The two resulting South African genomes were treated as candidate *B. cereus s.l.* genomes.

Whole-genome sequencing and assembly of *B. cereus s.l.* genomes from the Pennsylvania State University culture collection.

The genomes of 73 *B. cereus s.l.* strains from the Pennsylvania State University culture collection were additionally sequenced using a hybrid approach, which combined Illumina short-read sequencing with long-read sequencing. To generate Illumina reads, total genomic DNA was extracted from isolates using the E.Z.N.A Bacterial DNA extraction kit (Omega Bio-Tek, GA, USA) by following the manufacturer’s instructions. Extracted DNA was examined for quality and quantity using Nanodrop One (Thermo Fisher Scientific, MA, USA) and Qubit 3 (Thermo Fisher Scientific, MA, USA), respectively. DNA libraries were prepared using a NexteraXT library preparation kit (Illumina, CA, USA). Samples were sequenced using an Illumina NextSeq with 150 bp paired-end reads. Illumina adapters and low-quality reads were trimmed using Trimmomatic v0.39 (Supplementary Text). Trimmed read qualities were assessed using FastQC v0.11.9 (default settings).

To generate long reads, DNA was extracted using the QIAamp DNA blood mini kit (Qiagen, Hilden, Germany) by following the manufacturer’s instructions with additional steps for cell lysis. Briefly, 3 loopfuls of biomass grown at 30°C on Brain-Heart Infusion (BHI) agar (BD Biosciences, NJ, USA) were collected and mixed with a phosphate-buffered saline (PBS) buffer (137 mM NaCl, 1.7 mM KCl, 10 mM Na_2_HPO_4_, 1.8 mM KH_2_PO_4_). One gram of 0.1 mm zirconia/silica beads (Biospec Products, OK, USA) were added to tubes, which were then vortexed in a horizontal position to disrupt cell walls. The Oxford Nanopore Technologies (ONT) RBK-004 rapid barcoding kit was used for the library preparation by following manufacturer’s instructions, and 4 libraries were pooled for sequencing (ONT, Oxford, UK). An R9 flow cell (FLO-MIN106; ONT, Oxford, UK) was used for sequencing using a MinION Mk1C device (ONT). Raw sequencing signal files were used for high accuracy basecalling and adapter trimming using Guppy v6.0.1, and low-quality reads were trimmed using FiltLong v0.2.1 (Supplementary Text). The quality of trimmed reads was assessed using FastQC v0.11.9 (default settings). Trimmed Illumina short reads and trimmed ONT long reads were used for hybrid genome assembly with Unicycler v0.5.0 (using default parameters). The 73 resulting genomes were treated as candidate *B. cereus s.l.* genomes.

### Evaluation of genome quality

Overall, a total of 6,901 candidate *B. cereus s.l.* genomes were sequenced, assembled, and/or aggregated for this study: (i) 6,826 genomes derived from publicly available data (3,428 previously assembled, publicly available GenBank genomes from NCBI’s Assembly database, and 3,398 genomes assembled here from publicly available sequencing reads), (ii) two genomes derived from South African strains sequenced in this study, and (iii) 73 genomes derived from strains in the Pennsylvania State University culture collection, which were sequenced in this study.

The quality of each assembled genome was evaluated using QUAST v5.0.2^49^ (default settings, except “--min-contig 1”) and CheckM v1.1.3^50^ (using the “lineage_wf” workflow and default settings). Assembled genomes with (i) N50 < 20k (via QUAST), (ii) completeness < 95% (via CheckM), and (iii) contamination > 5% (via CheckM) were flagged as poor quality (*n* = 6,001 high-quality, candidate *B. cereus s.l.* genomes).

### Taxonomic assignment and sequence typing

Each assembled genome (*n* = 6,901; see section “Evaluation of genome quality” above) was supplied as input to BTyper3 v3.3.3,^20^ which was used to perform the following analyses: (i) taxonomic assignment using a standardized, strain-specific genomospecies-subspecies-biovar (GSB) taxonomic framework for *B. cereus s.l.* (the 2020 *B. cereus s.l.* GSB taxonomy);^3,10,20^ (ii) calculation of average nucleotide identity (ANI) values relative to the type strain genomes of all validly published and effective *B. cereus s.l.* species via FastANI^51^ (*n* = 28 species, accessed 8 August 2022; see section “Acquisition and assembly of putative *B. cereus s.l.* genomes from publicly available genomic data” above); (iii) ANI-based *B. cereus s.l.* pseudo-gene flow unit assignment;^20,52^ (iv) *in silico* multi-locus sequence typing (MLST) using PubMLST’s seven-gene MLST scheme for “*B. cereus*” (accessed 12 August 2022);^53^ (v) *in silico panC* phylogenetic group assignment^33^ using BTyper3’s adjusted eight-group *panC* assignment framework;^20^ (vi) detection of selected *B. cereus s.l.* virulence factors using translated nucleotide BLAST^54^ and BTyper3’s default settings (Supplementary Text)^31^; (vii) detection of insecticidal toxin-encoding genes using BTyper3’s default settings (Supplementary Text).^20^

Each assembled genome additionally underwent taxonomic assignment using the Genome Taxonomy Database (GTDB) Toolkit (GTDB-Tk) v2.1.0 and GTDB-Tk reference database version r207 (release207_v2).^55,56^ Genomes that (i) were flagged as poor quality (see section “Evaluation of genome quality” above) and/or (ii) did not belong to GTDB’s “Bacillus_A” genus were excluded from subsequent analyses. Overall, the sequencing, genome assembly, and genome aggregation efforts conducted here resulted in a final set of 5,976 high-quality *B. cereus s.l.* genomes, which were used to construct the initial BTyperDB database (Supplementary Table S1).

### Metadata acquisition and curation

Strain metadata (e.g., isolation source, isolation year, geographic location of isolation) associated with the 75 genomes sequenced in this study were provided by collaborators (Supplementary Table S1). Metadata associated with genomes aggregated and/or assembled from publicly available sequencing data were acquired from NCBI’s BioSample database. Briefly, for each of the 5,901 genomes derived from publicly available data, a text file containing the genome’s metadata was downloaded from NCBI’s BioSample database using the rentrez v1.2.3 package^57^ in R v3.6.3,^58^ using the genome’s BioSample accession as the query (Supplementary Table S1).

For each of the 5,976 high-quality *B. cereus s.l.* genomes, curators assigned (i) a year of isolation, (ii) a geographic location of isolation, and (iii) an isolation source by manually examining the genome’s (a) BioSample record, (b) BioProject record, and (c) publication(s) linked to the BioProject record. If any of this information was not available, (d) strain names were queried in Google with the goal of identifying the genome in an unlinked publication and/or external database (Supplementary Text).

Independent of their isolation source assignments, genomes were further classified based on the following: (a) whether the associated strain was known to be responsible for a human illness/infection; (b) the type of human illness/infection the associated strain was reportedly responsible for; (c) whether the associated strain was known to be responsible for >1 human illness cases (e.g., an outbreak; Supplementary Text). If the strain was responsible for an outbreak or cluster of illnesses, a unique identifier was created for the outbreak/cluster, so that all strains isolated in conjunction with the same event could be grouped together. For example, 29 genomes of *B. cereus s.l.* strains isolated in conjunction with an emetic outbreak linked to refried beans in New York state in 2016 were all given the identifier “FBOE:2016USANYSBEANS” (Supplementary Table S1, Supplementary Text). Figures showcasing BTyperDB metadata were constructed in R (Supplementary Text).

### Database and web portal construction

To construct the BTyperDB database, two input CSV files were created: (i) one file that describes the column types (e.g., text or number, and whether it should be displayed), (ii) one file that contains all table content. The table content file (ii) was converted into an SQLite3 database for fast querying. The portal back-end provides a RESTful API, implemented using Flask in Python. The front-end was implemented using React, with a map display implemented using the Highcharts library. Nginx was used to direct URLs to the front- and back-end, respectively.

### Species-level phylogeny construction

To construct a maximum likelihood phylogeny of all 5,976 *B. cereus s.l.* genomes in BTyperDB, Prokka v1.14.6^59^ was used to identify and annotate genes in each genome (using default settings and the “Bacteria” kingdom database). GFF files produced by Prokka were supplied as input to Panaroo v1.2.7,^60^ which was used to construct a core genome alignment of all 5,976 *B. cereus s.l.* genomes, plus an outgroup genome (Supplementary Text). IQ-TREE v2.2.0.3^61^ was used to construct a maximum likelihood phylogeny, using an alignment of core SNPs detected in all 5,976 genomes as input (Supplementary Text). The resulting *B. cereus s.l.* phylogeny was displayed and annotated in iTOL v6,^62^ using metadata from BTyperDB.

All aforementioned steps were repeated to construct species-level phylogenies for GTDB’s (i) *B. tropicus*, (ii) *B. cereus*, and (iii) *B. nitratireducens* species (Supplementary Table S1, Supplementary Text). Core SNPs produced by Panaroo and snp-sites were additionally supplied to RhierBAPS v1.1.4,^63^ which was used to partition each species into clusters using default settings and two clustering levels.

### Pairwise ANI calculations

ANI values were calculated between all genomes assigned to GTDB’s (i) *B. tropicus*, (ii) *B. cereus*, and (iii) *B. nitratireducens* species using FastANI v1.32 (Supplementary Table S1).^51^ Pairwise ANI values were displayed in a heatmap using the pheatmap v1.0.12 R package (https://cran.r-project.org/web/packages/pheatmap/; Supplementary Text). For *B. tropicus*, a two-dimensional graph of pairwise ANI values was additionally constructed using the “ANI.graph” function in the bactaxR v0.2.2^10^ R package (Supplementary Text).

### Within-lineage variant calling and phylogeny construction

Snippy v4.6.0 (https://github.com/tseemann/snippy) was used to identify core SNPs within each of the following lineages (selected via holistic consideration of MLST, pairwise ANI values, RhierBAPS clustering, and species-level phylogenomic topology): (i) the ST78 clade, (ii) ST234, and (iii) ST932. After alignment cleaning, recombination removal,^64^ and SNP filtering, IQ-TREE was used to construct a maximum likelihood phylogeny for each lineage (Supplementary Text).

## Data availability

All assembled genomes and metadata generated and curated in this study are publicly available and can be downloaded via BTyperDB (www.btyper.app).

## Code availability

The BTyperDB source code is open-source and available at https://github.com/henriksson-lab/btyper_website. Additional analyses performed in the study are described in the manuscript text and/or Supplementary Text.

## ACKNOWLEDGMENTS

V Ramnath, V Raghuram, HG, and LMC were supported by the SciLifeLab & Wallenberg Data Driven Life Science Program (grant: KAW 2020.0239). This research was conducted using the resources of High Performance Computing Center North (HPC2N) and the Uppsala Multidisciplinary Center for Advanced Computational Science (UPPMAX). Specifically, the computations/data handling were enabled by resources in projects (i) hpc2n2023-117 and hpc2nstor2023-041, and (ii) SNIC 2022/22-1160, SNIC 2022/23-600, and NAISS 2023/6-285 provided by the National Academic Infrastructure for Supercomputing in Sweden (NAISS) at HPC2N and UPPMAX, respectively, partially funded by the Swedish Research Council through grant agreement no. 2022-06725. Additional computational resources were provided by the European Molecular Biology Laboratory (EMBL) HPC and BEAGLE-COMPUTE (Umeå University). Fig. 1 was created with BioRender (https://www.biorender.com/).

## AUTHOR CONTRIBUTIONS

JH and V Ramnath were responsible for BTyperDB front-end development, while ML, V Ramnath, and LMC were responsible for back-end development. Computational analyses were performed by LMC and V Ramnath, with help from TC, V Raghuram, and HG. Metadata curation and standardization was led by PM and LMC with major contributions from AJB and ASH; additional curated metadata was provided by TC, JK, and IM. Data generation was performed by TC, XW, JK, RP, and IM. All authors assisted with data aggregation and results interpretation. LMC, V Ramnath, and JH wrote and prepared the manuscript with input from all authors. LMC conceived the study, which was funded by LMC and JH.

