## Supplementary Figure S1 for "A community-curated, global atlas of *Bacillus cereus sensu lato* genomes for epidemiological surveillance"

**A**

**Microbe sample from Bacillus cereus**

| | |
|---|---|
| Identifiers | BioSample: SAMN08688986; Sample name: FSL R9-6395; SRA: SRS3037277 |
| Organism | Bacillus cereus<br>cellular organisms; Bacteria; Terrabacteria group; Bacillota; Bacilli; Bacillales; Bacillaceae; Bacillus; Bacillus cereus group |
| Package | Microbe; version 1.0 |
| Attributes | **strain** FSL R9-6395<br>**isolation source** food<br>**collection date** 2016<br>**geographic location** USA: New York state<br>**sample type** Cell culture |
| BioProjects | PRJNA514245<br>Retrieve all samples from this project<br><br>PRJNA437714 Bacillus cereus<br>Retrieve all samples from this project |
| Submission | Cornell University, Laura Carroll; 2018-03-12 |

Accession: SAMN08688986 ID: 8688986
BioProject SRA

**B**

**Sample from Bacillus cereus AH187**

| | |
|---|---|
| Identifiers | BioSample: SAMN02604058; Sample name: CP001177 |
| Organism | Bacillus cereus AH187<br>cellular organisms; Bacteria; Terrabacteria group; Bacillota; Bacilli; Bacillales; Bacillaceae; Bacillus; Bacillus cereus group; Bacillus cereus |
| Attributes | **strain** AH187<br>**culture collection** DSM:4312 |
| BioProject | PRJNA17715 Bacillus cereus AH187<br>Retrieve all samples from this project |

Accession: SAMN02604058 ID: 2604058
BioProject Nucleotide

**Supplementary Figure S1.** Examples of NCBI BioSample records linked to *Bacillus cereus sensu lato* (*s.l.*) genomes. Panel (A) represents a typical but imperfect contemporary BioSample record. Useful metadata attributes, such as the year of isolation (2016), geographic location of isolation (New York state, United States of America), and isolation source (food) are reported, and the genome's species is listed as *Bacillus cereus*. However, the isolation source (food) is vague; it is unclear which type of food this strain was isolated from, nor is it clear if the strain was responsible for a human illness or outbreak. A literature search reveals that this particular isolate originated from refried beans and was responsible for an emetic foodborne outbreak (doi: 10.3389/fmicb.2019.00144), meaning that much more information is available for this genome than what was provided by the submitter. Further, despite being listed as *Bacillus cereus*, users of the Genome Taxonomy Database (GTDB) taxonomy would consider this genome to be a member of *Bacillus paranthracis.* Finally, this genome is represented by Illumina short reads, which were deposited in NCBI's Sequence Reach Archive (SRA) database; no assembled genome is available, meaning that this genome has been excluded from previous large-scale genomic studies of *B. cereus s.l.* Panel (B) represents a typical BioSample record for a historical strain; strain AH187 is a well-known emetic strain isolated from the vomit of a person who had eaten cooked rice, which was linked to an emetic outbreak in 1972. However, this information is not available in the BioSample record and is thus not included in pathogen surveillance tools that rely on BioSample records alone for metadata.
