## Supplementary Figure S2 for "A community-curated, global atlas of *Bacillus cereus sensu lato* genomes for epidemiological surveillance"

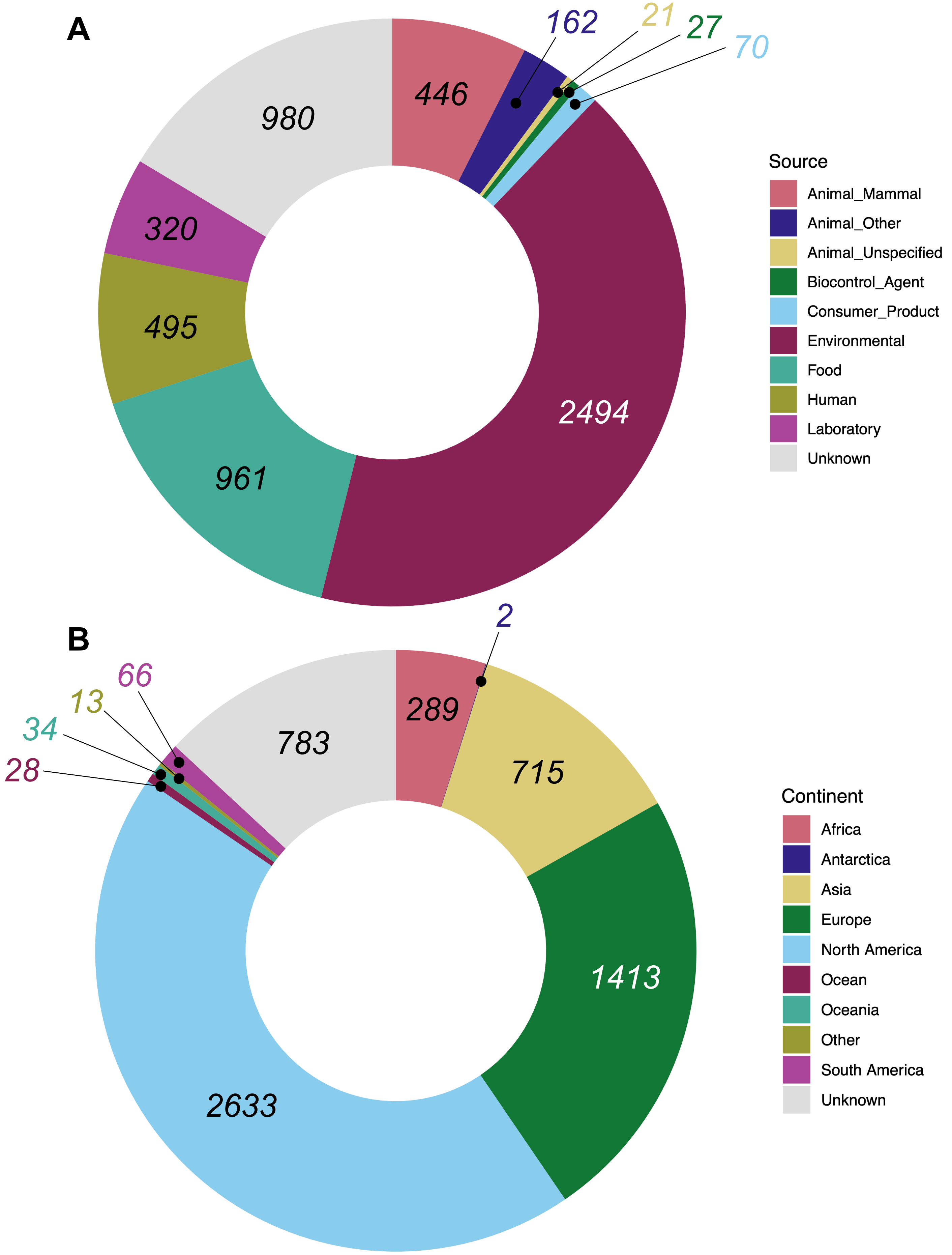


**Supplementary Figure S2.** Proportion of BTyperDB genomes isolated (A) from a given source (the BTyperDB Ontology "Source_1" attribute), or (B) from a given geographic location (the BTyperDB Ontology "Continent" attribute). Numeric labels denote the number of genomes assigned to a given category.
