## Supplementary Figure S3 for "A community-curated, global atlas of *Bacillus cereus sensu lato* genomes for epidemiological surveillance"

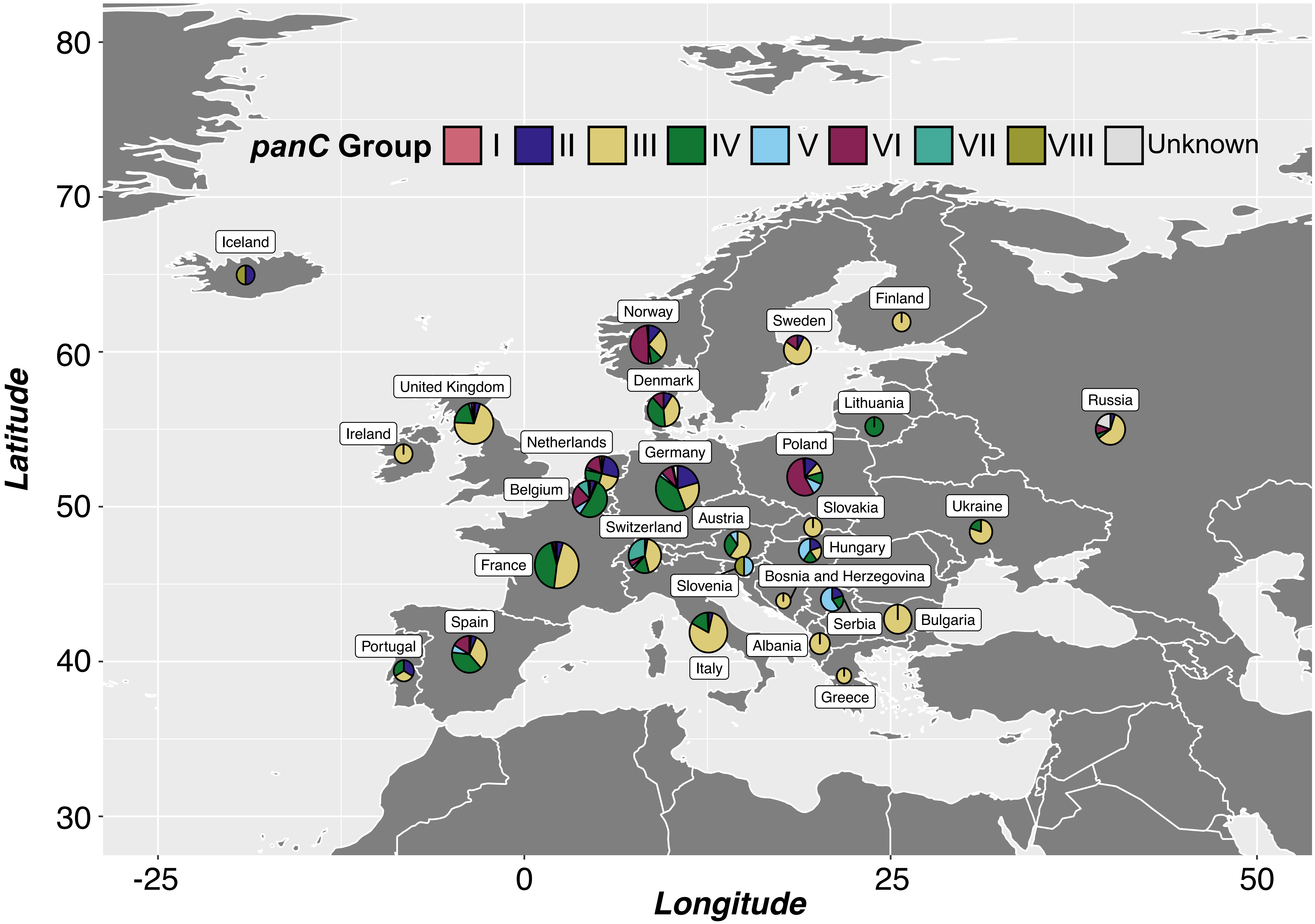


Supplementary Figure S3. Country of origin for all B. cereus s.l. genomes in BTyperDB, which were reportedly isolated in "Europe" (as defined via pycountry). Pie charts are sized according to the number of genomes from a particular country and show the proportion of genomes assigned to each panC phylogenetic group (i.e., historically important "species"-like units delineated using the sequence of panC).
