## Supplementary Figure S4 for "A community-curated, global atlas of *Bacillus cereus sensu lato* genomes for epidemiological surveillance"

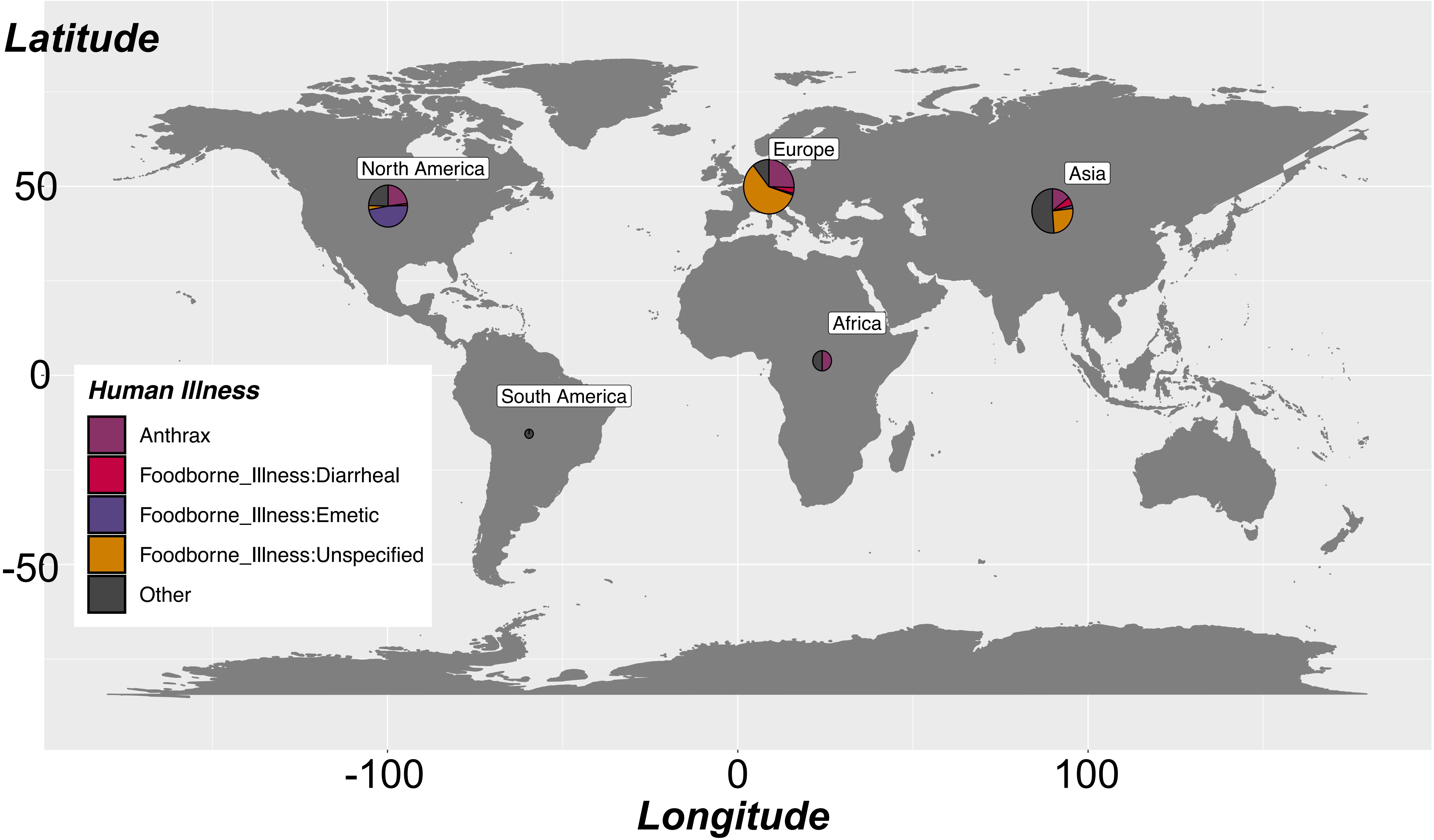


Supplementary Figure S4. Continent (as defined via pycountry) of origin for all B. cereus s.l. genomes in BTyperDB, which were linked to human illness cases. Pie charts are sized according to the number of human illness-linked genomes from a particular continent and show the proportion of genomes assigned to each illness type.
