## Supplementary Figure S5 for "A community-curated, global atlas of *Bacillus cereus sensu lato* genomes for epidemiological surveillance"

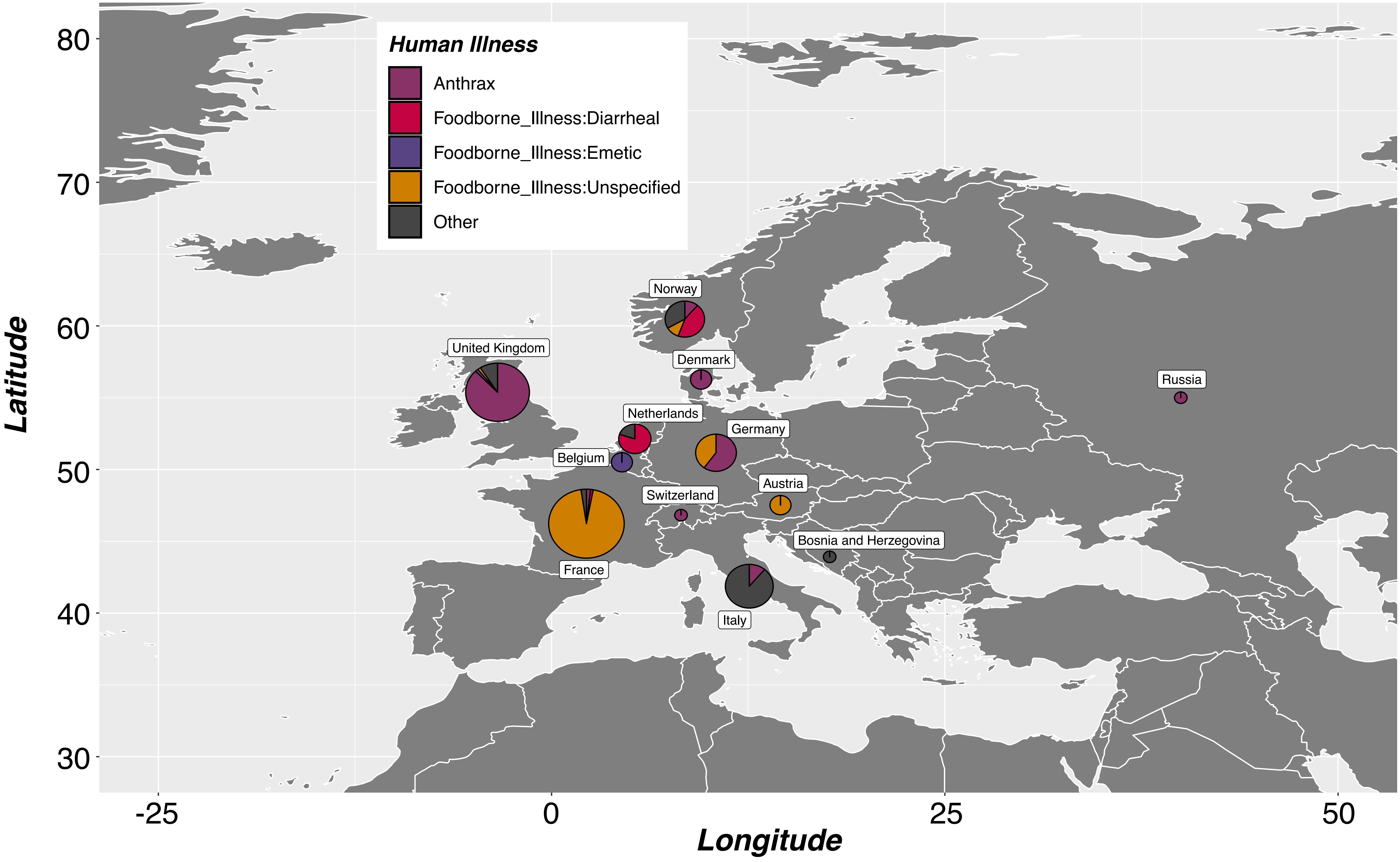


Supplementary Figure S5. Country of origin for all B. cereus s.l. genomes in BTyperDB, which were reportedly isolated in "Europe" (as defined via pycountry) and linked to human illness cases. Pie charts are sized according to the number of human illness-linked genomes from a particular country and show the proportion of genomes assigned to each illness type.
