## Supplementary Figure S6 for "A community-curated, global atlas of *Bacillus cereus sensu lato* genomes for epidemiological surveillance"

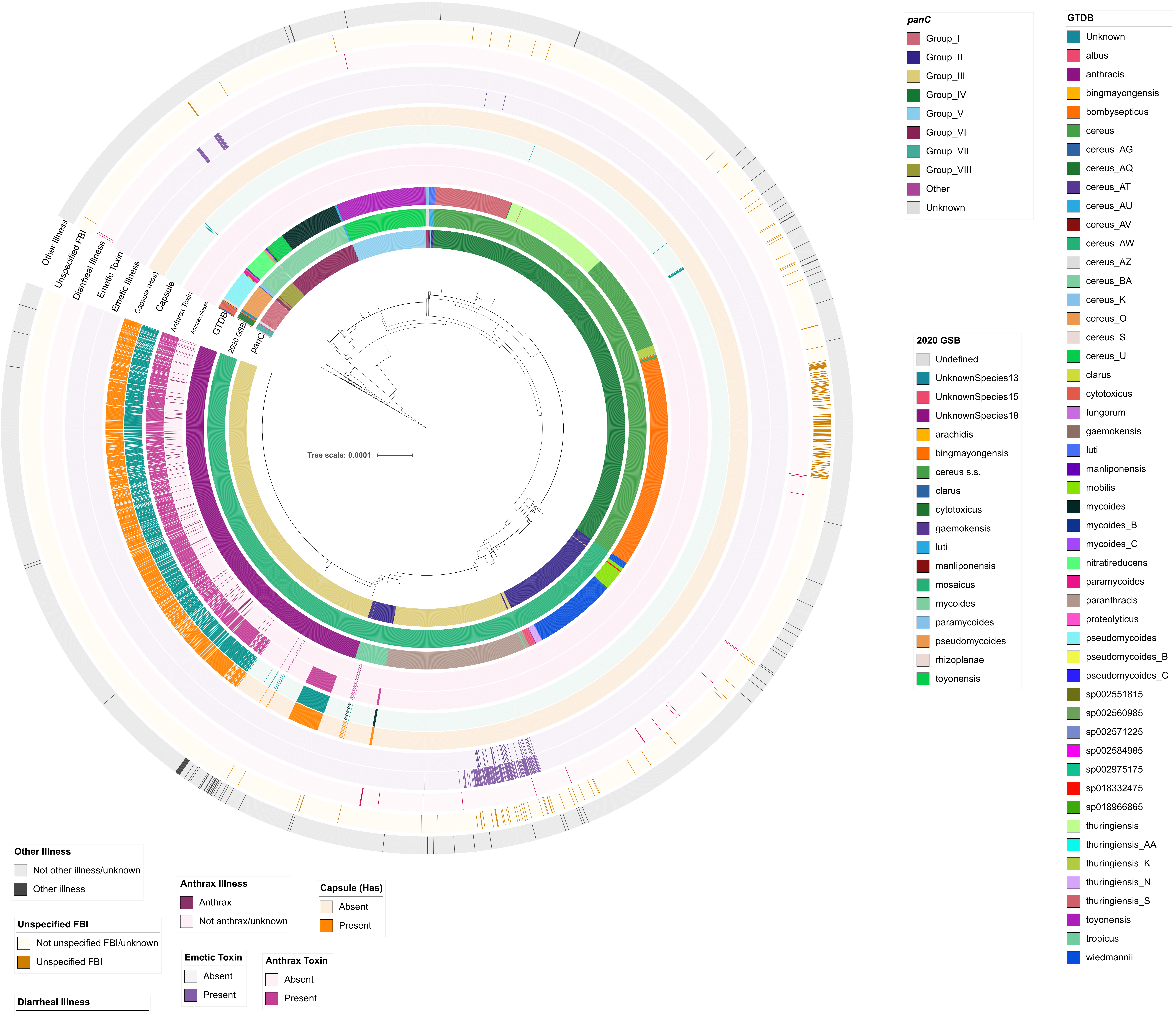

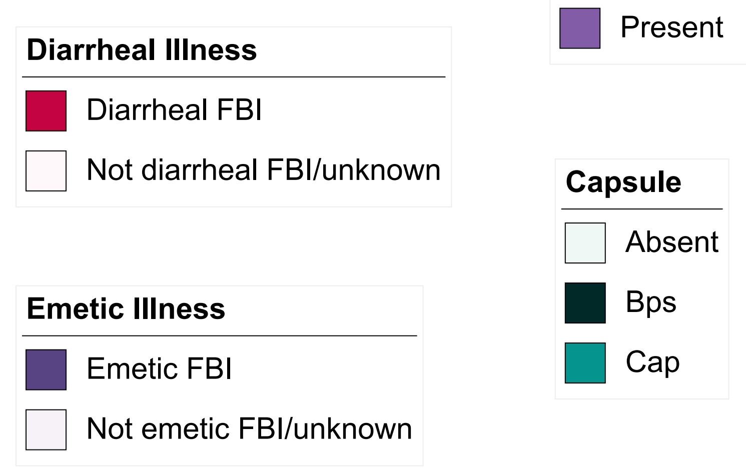


Supplementary Figure S6. Maximum likelihood phylogeny of 5,976 B. cereus s.l. genomes in BTyperDB. Color strips surrounding the phylogeny denote (from interior to exterior): (i) panC phylogenetic group (i.e., historically important “species”-like units delineated using the sequence of panC; “panC”); (ii) genomospecies assigned using the 2020 B. cereus s.l. Genomospecies-Subspecies-Biovar taxonomy (“2020 GSB”); (iii) genomospecies assigned using the Genome Taxonomy Database Toolkit (GTDB-Tk; “GTDB”); (iv) whether a genome was linked to a human anthrax case or not (“Anthrax Illness”); (v) presence and absence of two or more anthrax toxin genes (cya, lef, and/or pagA; “Anthrax Toxin”); (vi) presence and absence of select capsular genes (all of capBCADE or bpsXABCDEFGH; “Capsule”); (vii) presence and absence of one or more hyaluronic acid capsule genes (hasACB; “Capsule (Has)”); (viii) whether a genome was linked to a human foodborne emetic intoxication case or not (“Emetic Illness”); (ix) presence and absence of emetic toxin (cereulide) synthetase-encoding cesABCD (“Emetic Toxin”); whether a genome was linked to a (x) human diarrheal illness case, (xi) unspecified human foodborne illness (FBI) case, or (xii) other human illness/infection, or not (“Diarrheal Illness”, “Unspecified FBI’, and “Other Illness”, respectively). The phylogeny is rooted using an outgroup (omitted for readability), with branch lengths in substitutions per site.
