## Supplementary Figure S7 for "A community-curated, global atlas of *Bacillus cereus sensu lato* genomes for epidemiological surveillance"

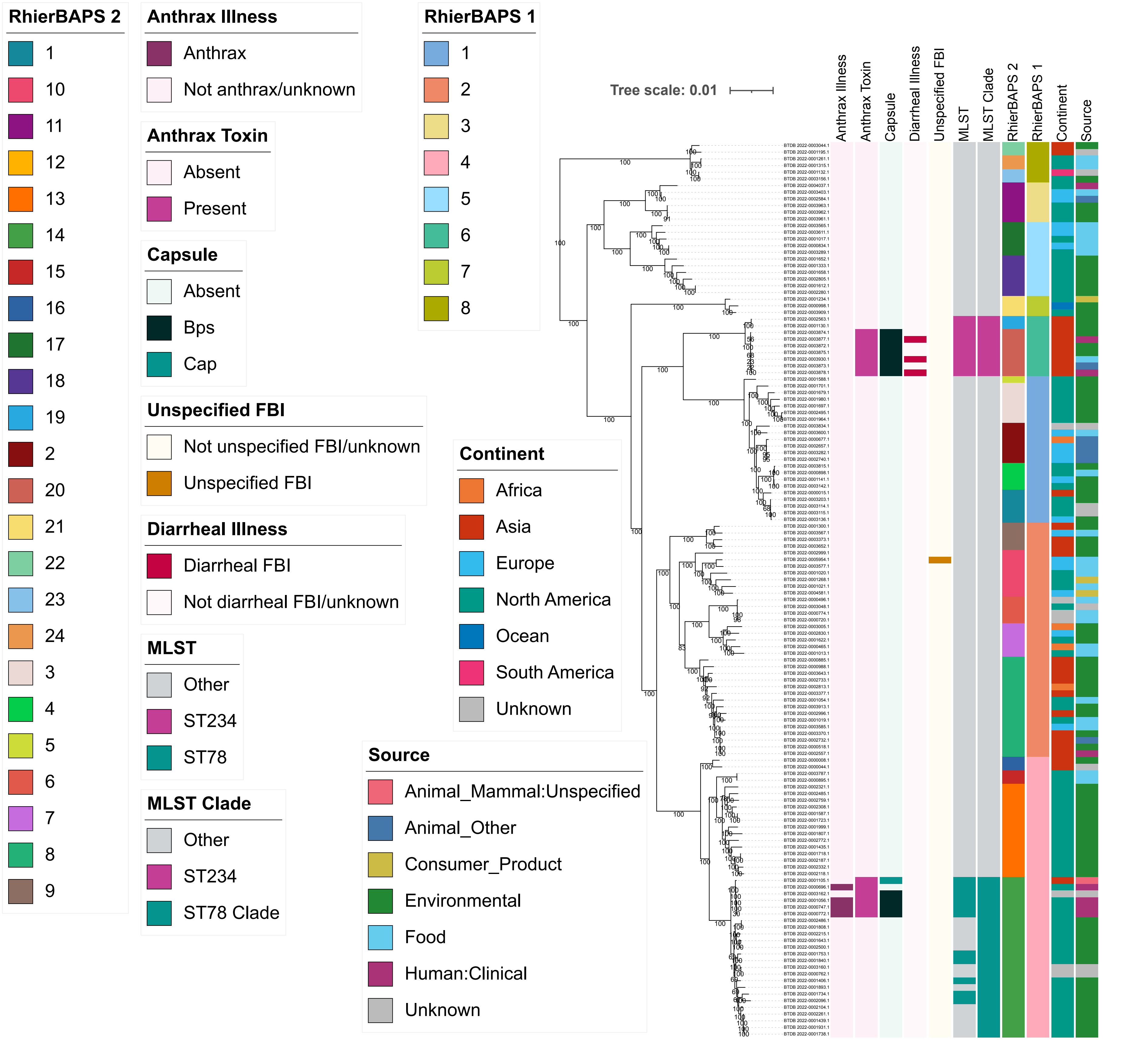


Supplementary Figure S7. Maximum likelihood phylogeny constructed using all genomes assigned to the B. tropicus species (per GTDB; n = 134). Color strips to the right of the phylogeny denote (from left to right): (i) whether a genome was linked to a human anthrax case or not; (ii) presence and absence of two or more anthrax toxin genes (cya, lef, and/or pagA); (iii) presence and absence of select capsular genes (all of capBCADE or bpsXABCDEFGH); (iv) whether a genome was linked to a human diarrheal illness case or not; (v) whether a genome was linked to an unspecified foodborne illness (FBI) case or not; (vi) sequence type, assigned using seven-gene multi-locus sequence typing (MLST); (vii) clade, assigned using MLST, average nucleotide identity (ANI) values, the B. tropicus phylogeny, and/or RhierBAPS; (viii) level 2 cluster assigned using RhierBAPS; (ix) level 1 cluster assigned using RhierBAPS; (x) continent and (xi) source from which each strain was reportedly isolated. Branch labels denote branch support percentages obtained using 1,000 replicates of the ultrafast bootstrap approximation. The phylogeny is rooted using an outgroup (omitted for readability), with branch lengths in substitutions per site.
