## Supplementary Figure S8 for "A community-curated, global atlas of *Bacillus cereus sensu lato* genomes for epidemiological surveillance"

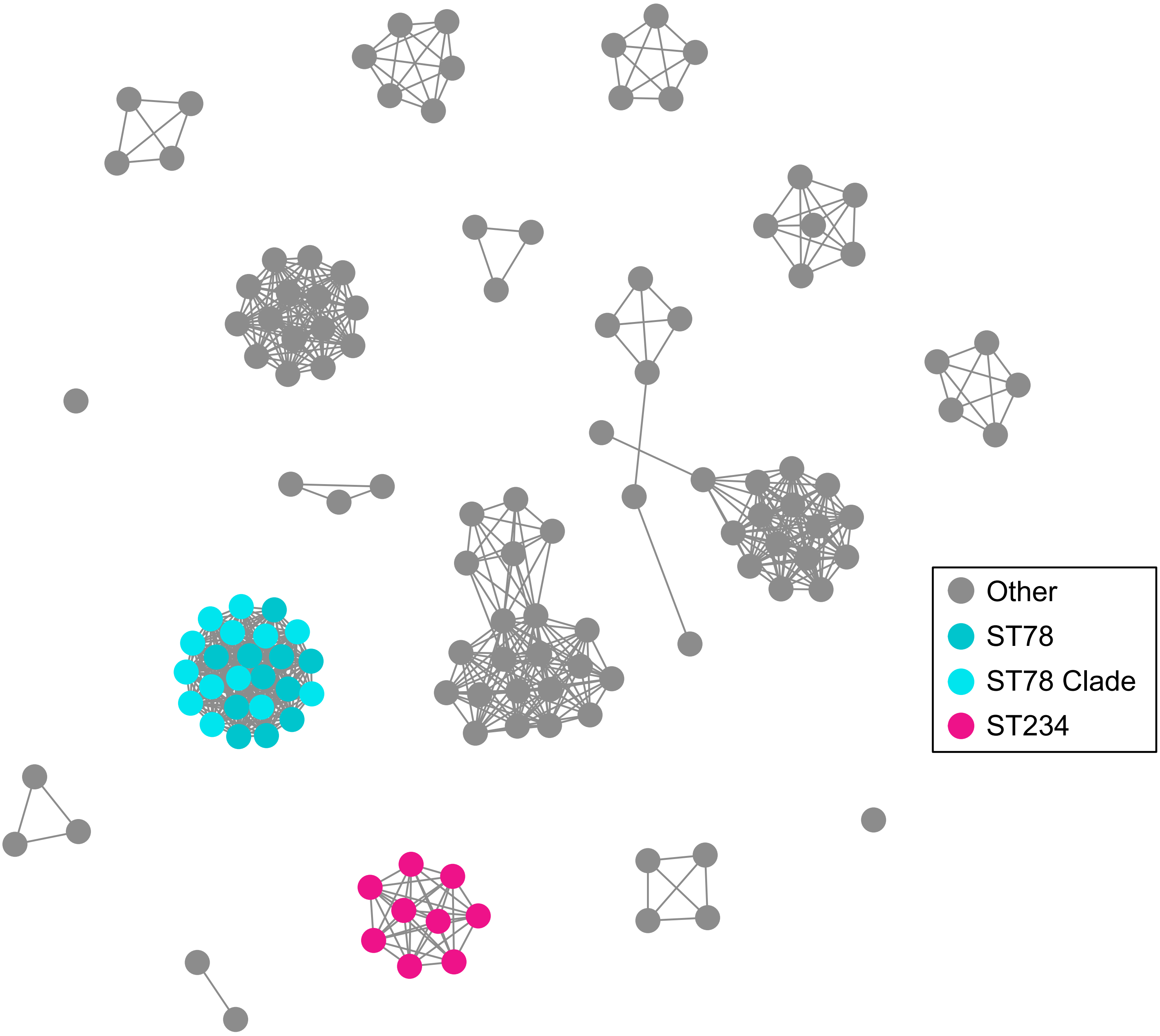


Supplementary Figure S8. Graph of pairwise average nucleotide identity (ANI) values calculated between all B. tropicus genomes (nodes; n = 17,956 pairwise comparisons between 134 genomes). Two genomes (nodes) are connected by an edge if they share ≥99 ANI with each other. Genomes (nodes) assigned to ST78 and/or the same “clade” as ST78 (based on the B. tropicus phylogeny, plus pairwise ANI values calculated between genomes; selected so that all ST78 genomes and ST78 clade genomes share ≥99 ANI with each other and <99 ANI with all other genomes) are colored blue (“ST78” and “ST78 Clade”). Genomes (nodes) assigned to ST234 are colored pink (“ST234”). Genomes (nodes) that did not belong to ST78, the ST78 clade, or ST234 are colored gray (“Other”). The graph was constructed using the ANI.graph function in the bactaxR R package (with “ANI_threshold = 99” and “graphout_niter = 1000000”). Pairwise ANI values were calculated using FastANI.
