## Supplementary Figure S9 for "A community-curated, global atlas of *Bacillus cereus sensu lato* genomes for epidemiological surveillance"

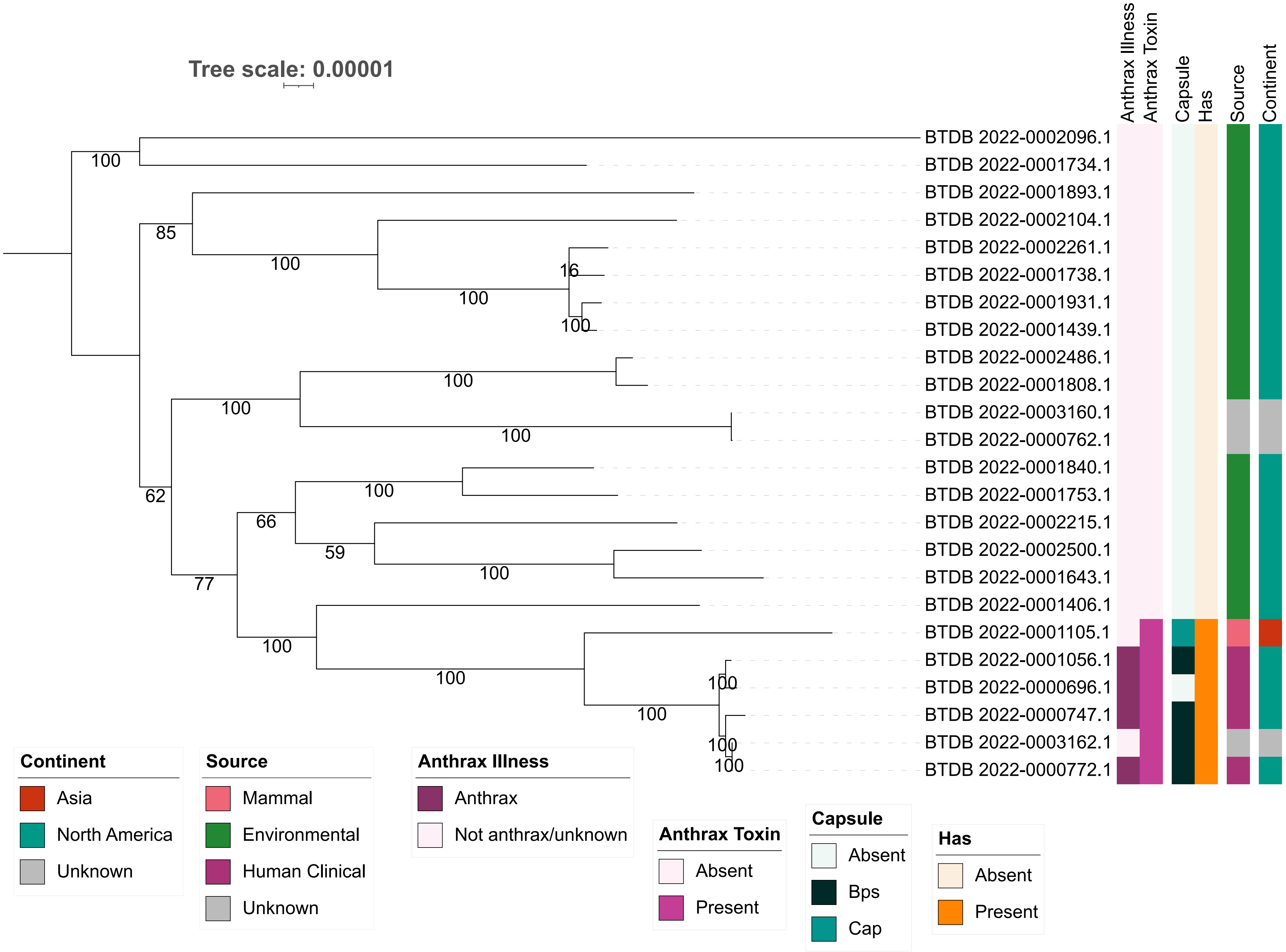


Supplementary Figure S9. Maximum likelihood phylogeny of all genomes assigned to the ST78 clade (n = 24). Color strips to the right of the phylogeny denote (from left to right): (i) whether a genome was linked to a human anthrax case or not; (ii) presence and absence of two or more anthrax toxin genes (cya, lef, and/or pagA); (iii) presence and absence of select capsular genes (all of capBCADE or bpsXABCDEFGH); (iv) presence and absence of one or more hyaluronic acid capsule-encoding genes (hasACB); (v) source and (vi) continent from which each strain was reportedly isolated. Branch labels denote branch support percentages obtained using 1,000 replicates of the ultrafast bootstrap approximation. The phylogeny is midpoint-rooted, with branch lengths in substitutions per site.
