## Supplementary Figure S10 for "A community-curated, global atlas of *Bacillus cereus sensu lato* genomes for epidemiological surveillance"

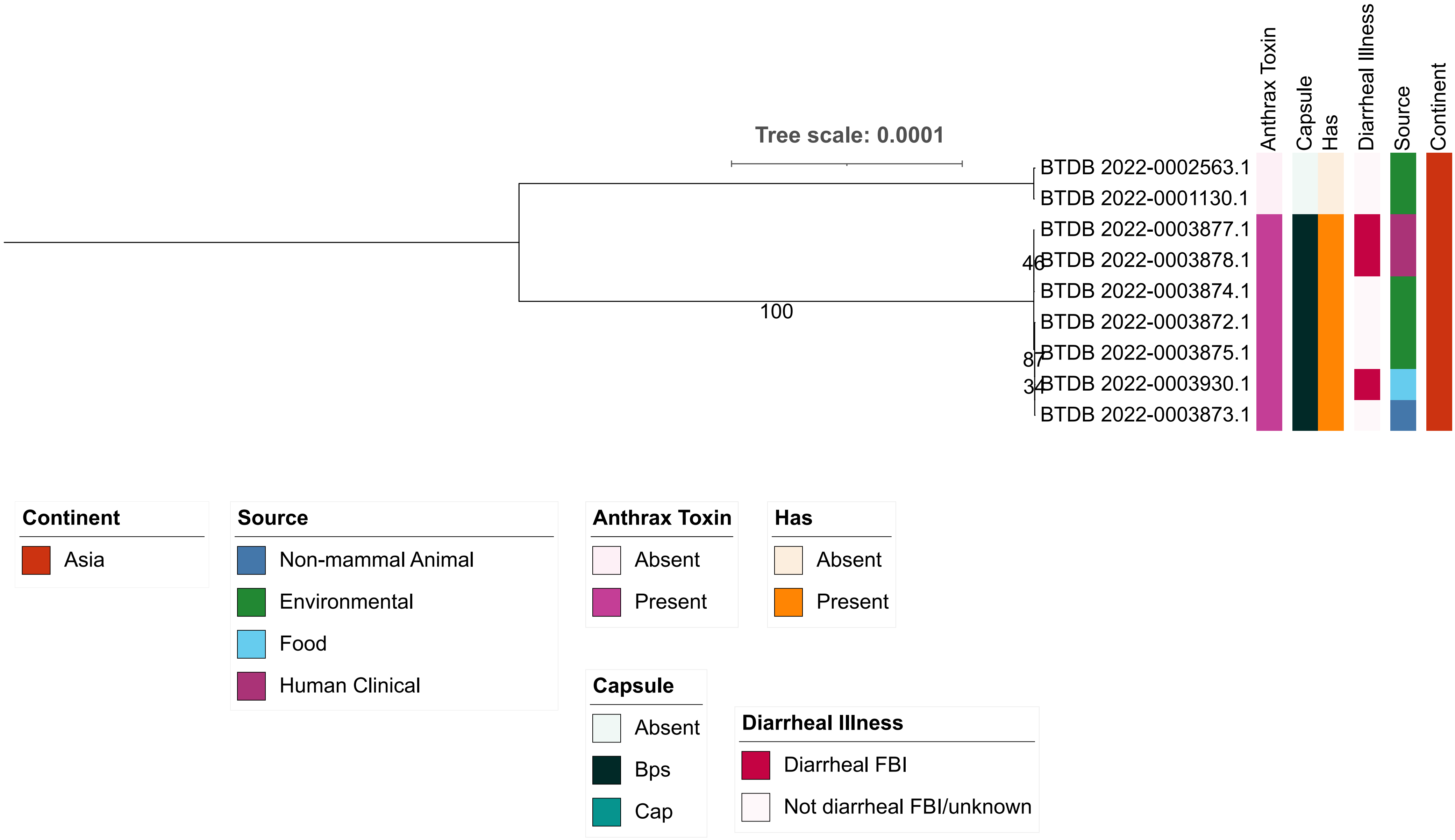


Supplementary Figure S10. Maximum likelihood phylogeny of all genomes assigned to ST234 (n = 9). Color strips to the right of the phylogeny denote (from left to right): (i) presence and absence of two or more anthrax toxin genes (cya, lef, and/or pagA); (ii) presence and absence of select capsular genes (all of capBCADE or bpsXABCDEFGH); (iii) presence and absence of one or more hyaluronic acid capsule-encoding genes (hasACB); (iv) whether a genome was linked to a human diarrheal illness case or not; (v) source and (vi) continent from which each strain was reportedly isolated. Branch labels denote branch support percentages obtained using 1,000 replicates of the ultrafast bootstrap approximation. The phylogeny is midpoint-rooted, with branch lengths in substitutions per site.
