## Supplementary Figure S11 for "A community-curated, global atlas of *Bacillus cereus sensu lato* genomes for epidemiological surveillance"

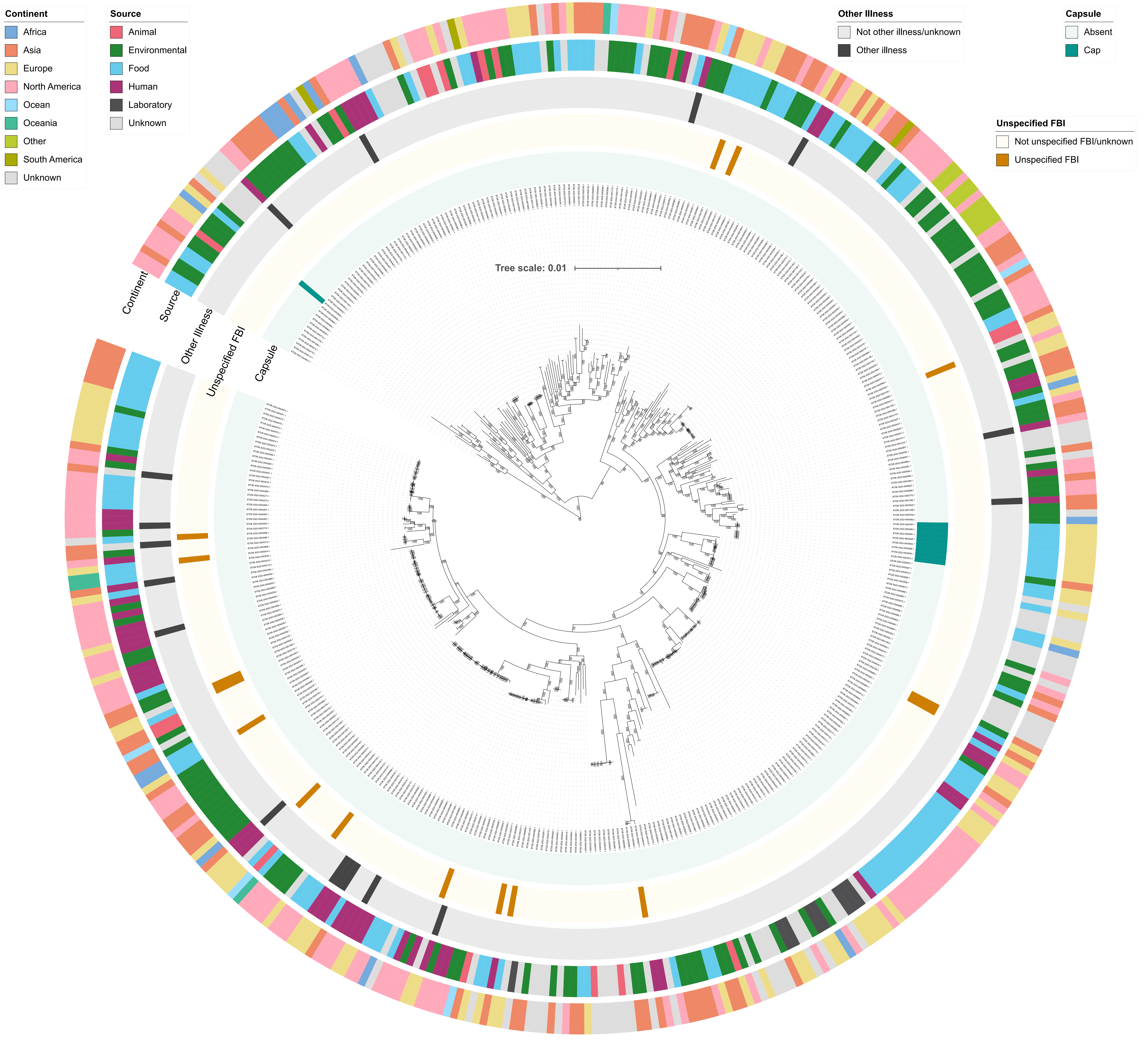


Supplementary Figure S11. Maximum likelihood phylogeny of all genomes assigned to GTDB's "B. cereus" species (n = 418). Color strips surrounding the phylogeny denote (from interior to exterior): (i) presence and absence of polyglutamate capsule-encoding genes (all of capBCADE; "Capsule"); whether a genome was linked to (ii) an unspecified human foodborne illness case ("Unspecified FBI"), or (iii) other human illness/infection, or not ("Other Illness"); the (iv) source and (v) continent from which each strain was reportedly isolated ("Source" and "Continent", respectively). Branch labels denote branch support percentages obtained using 1,000 replicates of the ultrafast bootstrap approximation. The phylogeny is rooted using an outgroup (omitted for readability), with branch lengths in substitutions per site. FBI, foodborne illness.
