## Supplementary Figure S12 for "A community-curated, global atlas of *Bacillus cereus sensu lato* genomes for epidemiological surveillance"

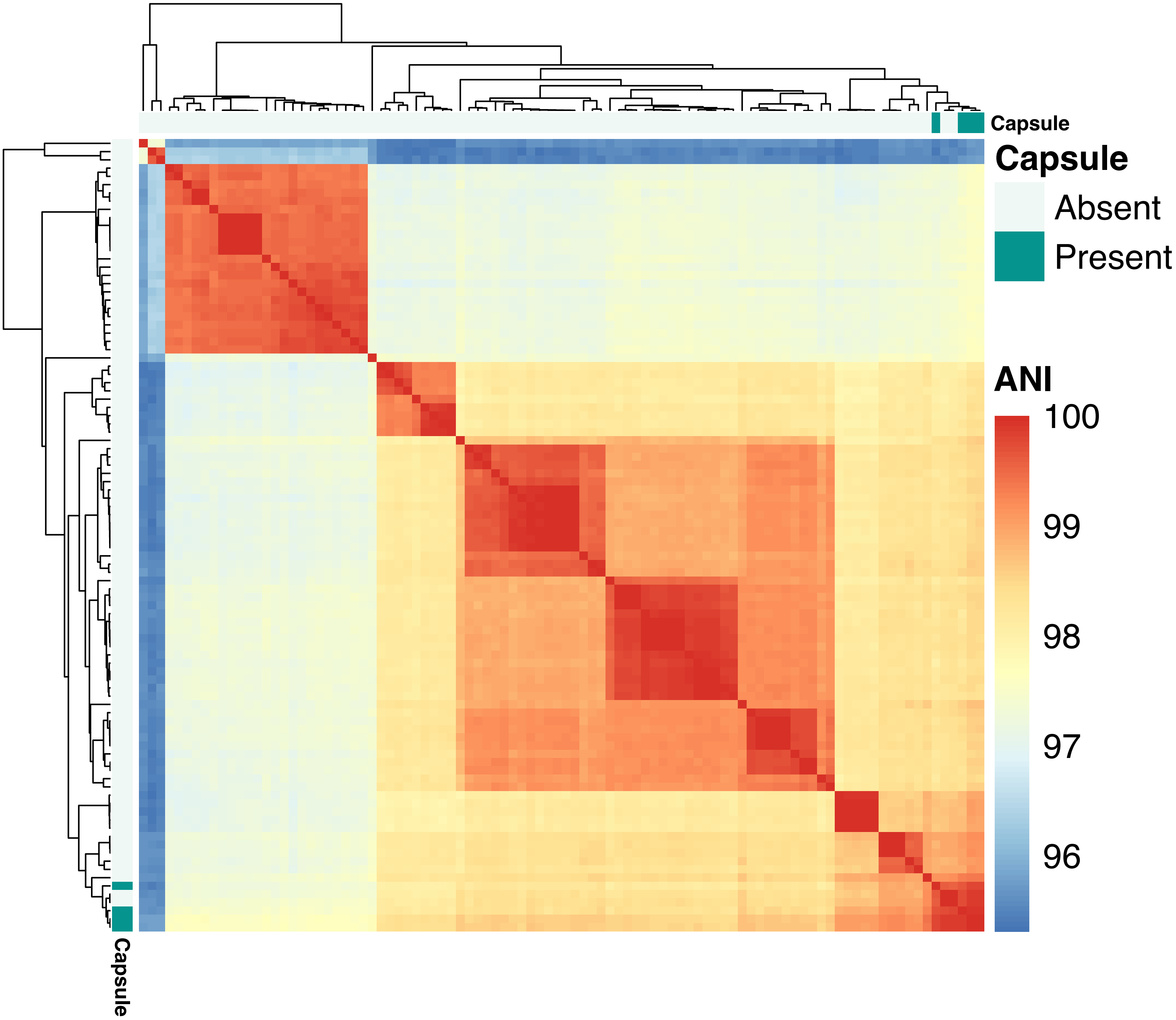


Supplementary Figure S12. Heatmap showcasing pairwise average nucleotide identity (ANI) values calculated between all GTDB B. nitratireducens genomes (n = 9,216 pairwise comparisons between 96 genomes). Presence and absence of polyglutamate capsule-encoding capBCADE in each genome are displayed in the color strips surrounding the heatmap. Dendrograms were constructed using average linkage hierarchical clustering, using ANI dissimilarities as input (i.e., 100-ANI). Pairwise ANI values were calculated using FastANI.
