## Supplementary Figure S13 for "A community-curated, global atlas of *Bacillus cereus sensu lato* genomes for epidemiological surveillance"

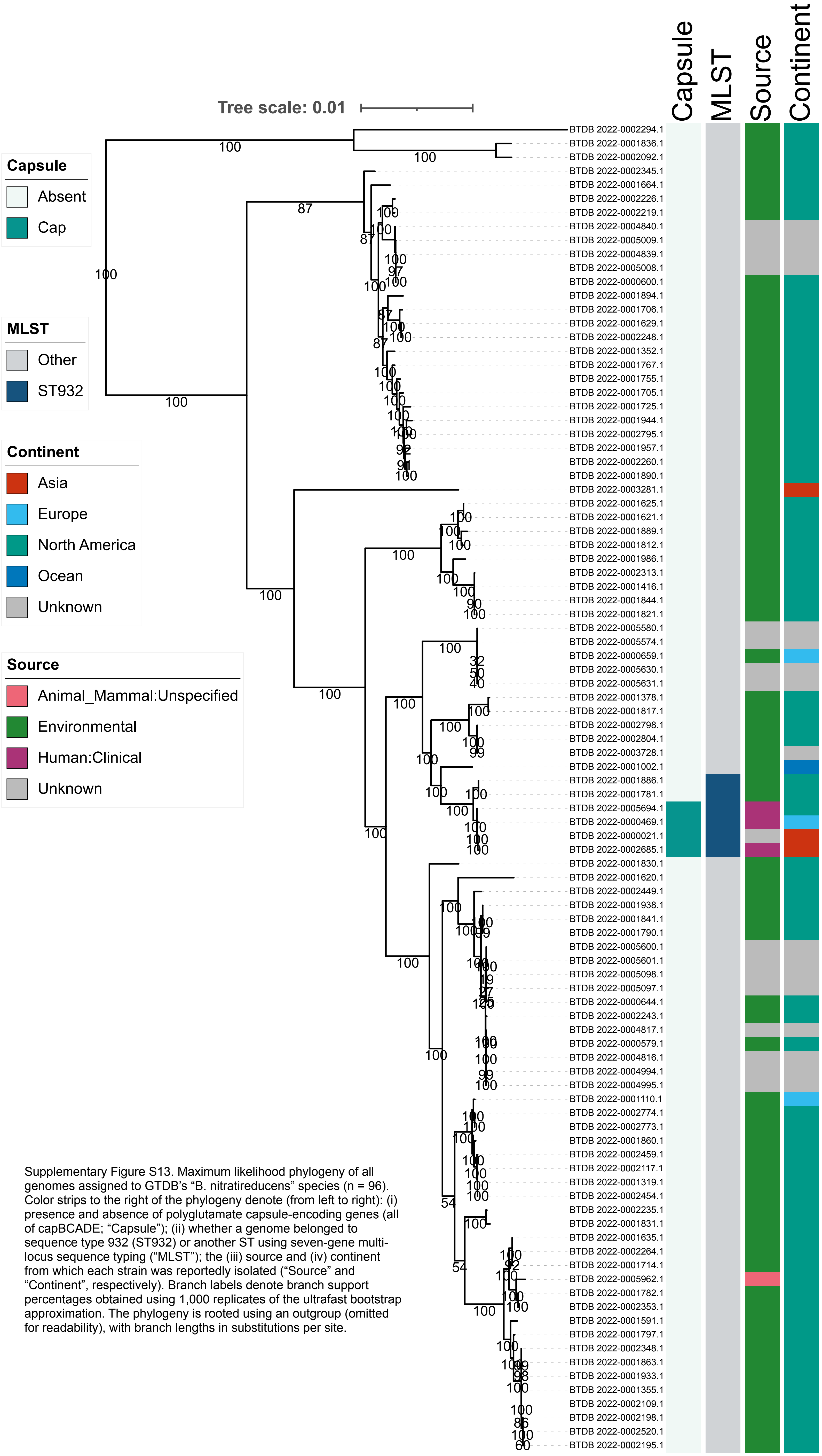


Supplementary Figure S13. Maximum likelihood phylogeny of all genomes assigned to GTDB's "B. nitratireducens" species (n = 96). Color strips to the right of the phylogeny denote (from left to right): (i) presence and absence of polyglutamate capsule-encoding genes (all of capBCADE; "Capsule"); (ii) whether a genome belonged to sequence type 932 (ST932) or another ST using seven-gene multi-locus sequence typing ("MLST"); the (iii) source and (iv) continent from which each strain was reportedly isolated ("Source" and "Continent", respectively). Branch labels denote branch support percentages obtained using 1,000 replicates of the ultrafast bootstrap approximation. The phylogeny is rooted using an outgroup (omitted for readability), with branch lengths in substitutions per site.
