## Supplementary Figure S14 for "A community-curated, global atlas of *Bacillus cereus sensu lato* genomes for epidemiological surveillance"

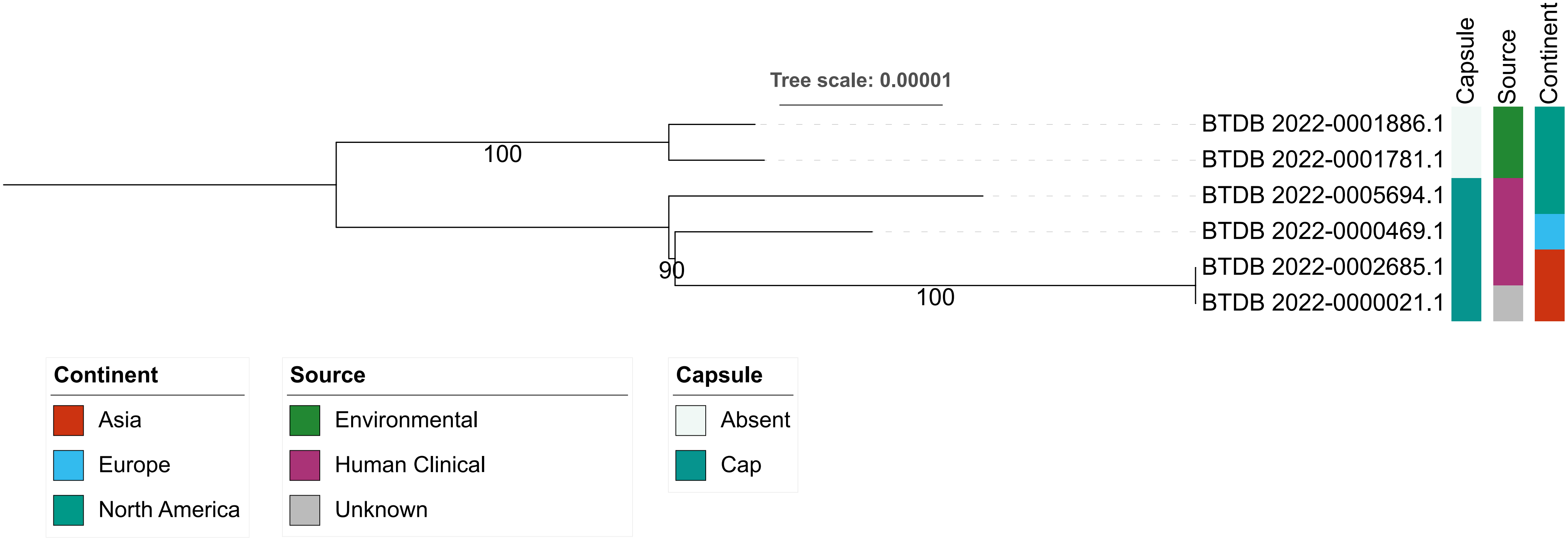


Supplementary Figure S14. Maximum likelihood phylogeny of all genomes assigned to ST932 (n = 6). Color strips to the right of the phylogeny denote (from left to right): (i) presence and absence of polyglutamate capsule genes (all of capBCADE); (ii) source and (iii) continent from which each strain was reportedly isolated. Branch labels denote branch support percentages obtained using 1,000 replicates of the ultrafast bootstrap approximation. The phylogeny is midpoint-rooted, with branch lengths in substitutions per site.
