## Supplementary Text for "A community-curated, global atlas of *Bacillus cereus sensu lato* genomes for epidemiological surveillance"

**Supplementary Text.** Supplementary methods for “A community-curated, global atlas of *Bacillus cereus sensu lato* genomes for epidemiological surveillance”.

***Supplementary Text***

**Acquisition and assembly of putative *B. cereus s.l.* genomes from publicly available genomic data.** All child nodes of the National Center for Biotechnology Information (NCBI) "*Bacillus cereus* group" taxonomic level (NCBI Taxonomy ID 86661) were treated as potential *B. cereus s.l.* members, and a list of their NCBI Taxonomy IDs was compiled (https://www.ncbi.nlm.nih.gov/taxonomy/?term=txid86661[Subtree], accessed 8 August 2022).[1] To ensure that no validly published and effective *B. cereus s.l.* species were absent from the list of *B. cereus s.l.* Taxonomy IDs prepared by NCBI, each of the following validly published and effective *B. cereus s.l.* species names were queried and manually added if not originally present in NCBI's *B. cereus s.l.* list ($n$ = 28 species, accessed 8 August 2022): *B. albus*,[2] *B. anthracis*,[3] "*B. arachidis*",[4] "*B. bingmayongensis*",[5] *B. cereus*,[6] "*B. clarus*",[7] *B. cytotoxicus*,[8] *B. fungorum*,[9] "*B. gaemokensis*",[10] *B. hominis*,[11] *B. luti*,[2] "*B. manliponensis*",[12] *B. mobilis*,[2] *B. mycoides*,[13] *B. nitratireducens*,[2] *B. pacificus*,[2] *B. paramobilis*,[11] *B. paramycoides*,[2] *B. paranthracis*,[2] *B. proteolyticus*,[2] *B. pseudomycoides*,[14] *B. rhizoplanae*,[15] *B. sanguinis*,[11] *B. thuringiensis*,[16] *B. toyonensis*,[17] *B. tropicus*,[2] *B. weihenstephanensis*,[18] *B. wiedmannii*.[19]

All assembled GenBank genomes available in NCBI's Assembly database (accessed 8 August 2022) were treated as candidate *B. cereus s.l.* genomes if the associated NCBI Taxonomy ID listed in the GenBank assembly summary file (https://ftp.ncbi.nlm.nih.gov/genomes/ASSEMBLY_REPORTS/assembly_summary_genbank.txt; accessed 8 August 2022) was included in the *B. cereus s.l.* Taxonomy ID list compiled here (Supplementary Table S2). All candidate *B. cereus s.l.* assembled genomes were downloaded from GenBank via NCBI's File Transfer Protocol (FTP) site (https://ftp.ncbi.nlm.nih.gov/; accessed 8 August 2022).

***Supplementary Text***

To search NCBI's Sequence Read Archive (SRA) database[20] for sequencing reads derived from potential *B. cereus s.l.* members, metadata in SRA was queried using Google Cloud's gcloud command line interface (CLI; accessed 8 August 2022). SRA accession numbers were considered to belong to potential *B. cereus s.l.* members if the value in SRA's "organism" field was equivalent to an organism name associated with one of the NCBI Taxonomy IDs compiled here (Supplementary Table S2). Additional search criteria included (i) the data was publicly available (consent = "public"), (ii) only genomic data was considered (assay_type = "WGS" and librarysource = "GENOMIC"), and (iii) the maximum number of data table rows was set to one billion (--max_rows 1000000000).

Using the compiled list of potential *B. cereus s.l.* genomes available in NCBI's (i) Assembly (GenBank) and (ii) SRA databases, a final list was compiled by taking the union of the NCBI BioSample[21] accession numbers associated with each genome or SRA run, respectively (accessed 8 August 2022). For BioSamples with an associated assembled genome, the assembled GenBank genome available in the Assembly database was used in subsequent steps. For BioSamples with no associated assembled genome (i.e., those with SRA data alone), FASTQ files associated with the BioSample were downloaded using the SRA Toolkit v2.11.0 "prefetch" and "fastq-dump" commands.[20] The quality of the resulting raw FASTQ files was assessed using FastQC v0.11.9[22] and MultiQC v1.12.[23] For SRA runs associated with paired-end Illumina reads, Shovill v1.1.0 (https://github.com/tseemann/shovill) was used to assemble each set of paired-end reads into contigs, using the following options: adapter trimming via Trimmomatic v0.39[24] enabled ("--trim"); a minimum contig length threshold of 200 bp

("--minlen 200"); a minimum contig coverage threshold of 10 ("--mincov 10"); SKESA v2.4.0[25] as the assembler ("--assembler skesa"). For SRA runs with single-end Illumina reads available, single-end reads were trimmed and assembled into contigs using Shovill's implementation of Trimmomatic and SKESA, respectively, as described above. For SRA runs with long reads available (i.e., PacBio or MinION), Unicycler v0.5.0[26] was used to assemble the genome using default settings with a minimum contig length of 200 bp ("--min_fasta_length 200"); when paired-end Illumina reads were additionally available, both long and short reads were supplied as input to Unicycler, and a hybrid assembly was constructed (Illumina reads were trimmed using Trimmomatic as described above). For SRA runs with paired-end Illumina and Roche 454 reads available, the paired-end Illumina reads were assembled into contigs using Shovill as described above, with 454 reads omitted.

The resulting paired-end Illumina reads associated with each strain were trimmed via Trimmomatic v0.38, using the following parameters: removal of Illumina adapters (“ILLUMINACLIP:TruSeq3-PE-2.fa:2:30:10:2:keepBothReads”), leading and trailing low quality or N bases (i.e., Phred quality < 3; “LEADING:3 TRAILING:3”), and reads < 36 bp in length (“MINLEN:36”). The quality of the resulting trimmed paired-end reads was assessed using FastQC, and Shovill was used to assemble each genome as described above, with the adapter trimming option omitted (see section "Acquisition and assembly of putative *B. cereus s.l.* genomes from publicly available genomic data" above). The two resulting South African genomes were treated as candidate *B. cereus s.l.* genomes.

**Whole-genome sequencing and assembly of *B. cereus s.l.* genomes from the Pennsylvania State University culture collection.** The genomes of 73 *B. cereus s.l.* strains from the Pennsylvania State University culture collection were additionally sequenced using a hybrid approach, which combined Illumina short-read sequencing with long-read sequencing. To generate Illumina reads, total genomic DNA was extracted from isolates using the E.Z.N.A Bacterial DNA extraction kit (Omega Bio-Tek, GA, USA) by following the manufacturer's instructions. Extracted DNA was examined for quality and quantity using Nanodrop One (Thermo Fisher Scientific, MA, USA) and Qubit 3 (Thermo Fisher Scientific, MA, USA), respectively. DNA libraries were prepared using a NexteraXT library preparation kit (Illumina, CA, USA). Samples were sequenced using an Illumina NextSeq with 150 bp paired-end reads. Illumina adapters and low-quality reads were trimmed using Trimmomatic v0.39 with a sliding window size of 4 and quality cutoff value of 15 (SLIDINGWINDOW:4:15). Reads shorter than 36 bp were excluded (MINLEN:36). Trimmed read qualities were assessed using FastQC v0.11.9.

To generate long reads, DNA was extracted using the QIAamp DNA blood mini kit (Qiagen, Hilden, Germany) by following the manufacturer's instructions with additional steps for cell lysis. Briefly, 3 loopfuls of biomass grown at 30°C on Brain-Heart Infusion (BHI) agar (BD Biosciences, NJ, USA) were collected and mixed with a phosphate-buffered saline (PBS) buffer (137 mM NaCl, 1.7 mM KCl, 10 mM

$Na2HPO4$, 1.8 mM $KH2PO4$). One gram of 0.1 mm zirconia/silica beads (Biospec Products, OK, USA) were added to tubes, which were then vortexed in a horizontal position to disrupt cell walls. The Oxford Nanopore Technologies (ONT) RBK-004 rapid barcoding kit was used for the library preparation by following manufacturer's instructions, and 4 libraries were pooled for sequencing (ONT, Oxford, UK). An R9 flow cell (FLO-MIN106; ONT, Oxford, UK) was used for sequencing using a MinION Mk1C device (ONT). Raw sequencing signal files were used for high accuracy basecalling and adapter trimming using Guppy v6.0.1 (https://community.nanoporetech.com/docs/prepare/library_prep_protocols/Guppy-protocol/v/gpb_2003_v1_revax_14dec2018/guppy-software-overview). Low-quality reads were trimmed using FiltLong v0.2.1 (https://github.com/rrwick/Filtlong), and the quality of trimmed reads was assessed using FastQC v0.11.9. Trimmed Illumina short reads and trimmed ONT long reads were used for hybrid genome assembly with Unicycler v0.5.0 (using default parameters). The 73 resulting genomes were treated as candidate *B. cereus s.l.* genomes.

**Evaluation of genome quality.** Overall, a total of 6,901 candidate *B. cereus s.l.* genomes were sequenced, assembled, and/or aggregated for this study: (i) 6,826 genomes derived from publicly available data (3,428 previously assembled, publicly available GenBank genomes from NCBI's Assembly database, and 3,398 genomes assembled here from publicly available sequencing reads), (ii) two genomes derived from South African strains sequenced in this study, and (iii) 73 genomes derived from strains in the Pennsylvania State University culture collection, which were sequenced in this study.

***Supplementary Text***

The quality of each assembled genome was evaluated using QUAST v5.0.2[28] (default settings, except "--min-contig 1") and CheckM v1.1.3[29] (using the "lineage_wf" workflow and default settings). Assembled genomes with (i) N50 < 20k (via QUAST), (ii) completeness < 95% (via CheckM), and (iii) contamination > 5% (via CheckM) were flagged as poor quality (*n* = 6,001 high-quality, candidate *B. cereus s.l.* genomes).

**Taxonomic assignment and sequence typing.** Each assembled genome (*n* = 6,901; see section "Evaluation of genome quality" above) was supplied as input to BTyper3 v3.3.3,[30] which was used to perform the following analyses: (i) taxonomic assignment using a standardized, strain-specific genomospecies-subspecies-biovar (GSB) taxonomic framework for *B. cereus s.l.* (the 2020 *B. cereus s.l.* GSB taxonomy);[30–32] (ii) calculation of average nucleotide identity (ANI) values relative to the type strain genomes of all validly published and effective *B. cereus s.l.* species via FastANI[33] (*n* = 28 species, accessed 8 August 2022; see section "Acquisition of *B. cereus s.l.* genomes and metadata" above); (iii) ANI-based *B. cereus s.l.* pseudo-gene flow unit assignment;[30,34] (iv) *in silico* multi-locus sequence typing (MLST) using PubMLST's seven-gene MLST scheme for "*B. cereus*" (accessed 12 August 2022);[35] (v) *in silico* *panC* phylogenetic group assignment[36] using BTyper3's adjusted eight-group *panC* assignment framework;[30] (vi) detection of selected *B. cereus s.l.* virulence factors using translated nucleotide BLAST[37] and BTyper3's default settings (i.e., with minimum identity and coverage thresholds of 70% and 80%, respectively)[31]; (vii) detection of insecticidal toxin-encoding genes using BTyper3's default settings (i.e., using translated nucleotide BLAST, with minimum identity and coverage thresholds of 50% and 70%, respectively).[30] In situations where virulence factor absence needed to be confirmed (e.g., to confirm the absence of anthrax toxin genes in some genomes), BTyper3's virulence factor detection workflow was run a second time, with minimum amino acid identity and coverage thresholds lowered to 40% and 0%, respectively.

Each assembled genome additionally underwent taxonomic assignment using the Genome Taxonomy Database (GTDB) Toolkit (GTDB-Tk) v2.1.0 and GTDB-Tk reference database version r207 (release207_v2).[38,39] Genomes that (i) were flagged as poor quality (see section "Evaluation of genome quality" above) and/or (ii) did not belong to GTDB's "Bacillus_A" genus were excluded from subsequent analyses. Overall, the sequencing, genome assembly, and genome aggregation efforts conducted here resulted in a final set of 5,976 high-quality *B. cereus s.l.* genomes, which were used to construct the initial BTyperDB database (Supplementary Table S1).

**Metadata acquisition and curation.** Strain metadata (e.g., isolation source, isolation year, geographic location of isolation) associated with the 75 genomes sequenced in this study were provided by collaborators (Supplementary Table S1). Metadata associated with genomes aggregated and/or assembled from publicly available sequencing data were acquired from NCBI's BioSample database. Briefly, for each of the 5,901 genomes derived from publicly available data, a text file containing the genome's metadata was downloaded from NCBI's BioSample database using the rentrez v1.2.3 package[40] in R v3.6.3,[41] using the genome's BioSample accession as the query (Supplementary Table S1).

Each of the 5,976 high-quality *B. cereus s.l.* genomes was assigned (i) a year of isolation, (ii) a geographic location of isolation, and (iii) an isolation source by manually examining the genome's (a) BioSample record, (b) BioProject record, and (c) publication(s) linked to the BioProject record. If any of this information was not available, (d) strain names were queried in Google with the goal of identifying the genome in an unlinked publication and/or external database. Metadata sources were recorded and categorized as “BioSample”, “BioProject”, “Publication”, or “URL” (if any metadata attribute was acquired from the genome’s BioSample record, BioProject record, one or more peer-reviewed publications, or one or more external websites, respectively); for metadata acquired from “Publication” or “URL” sources, the corresponding Digital Object Identifier (DOI) and Uniform Resource Locator (URL) were recorded, respectively.

For each genome’s geographic location of isolation, genomes were assigned a country of isolation using the pycountry v22.3.5 package (https://pypi.org/project/pycountry/) in Python v3.9.5 (https://www.python.org/). The assigned country was then given a continent designation using pycountry’s “country_alpha2_to_continent_code” function. In cases where more specific geographic information was available (e.g., a reported state, province, or city within a country), genomes were assigned a “region” name using pycountry’s principal subdivisions (i.e., subdivisions with no parent subdivision node). In cases where a principal subdivision could not be inferred, genomes were assigned a region of “Unknown”.

For each genome’s isolation source, curators first assigned each genome to one of the following principal source categories (termed “Source_1” in BTyperDB): (a) “Human” (the strain was isolated from a human); (b) “Animal_Mammal” (the strain was isolated from a mammal, excluding humans; also excluded animal/food products derived from mammals); (c) “Animal_Other” (the strain was isolated from an animal that was not a mammal; included insects and invertebrates); (d) “Animal_Unspecified” (the strain was isolated from an unknown, unspecified, or ambiguous animal); (e) “Food” (the strain was isolated directly from a food product, food ingredient, or dietary supplement with the potential for human oral consumption; excluded animal feed); (f) “Consumer_Product” (the strain was isolated from an item intended to be applied to/inside the human body, but was not intended for human oral consumption; e.g., nasal sprays, makeup, baby wipes); (g) “Biocontrol_Agent” (the strain was explicitly stated to be a biocontrol agent, biopesticide, and/or insecticide); (h) “Environmental” (the strain was isolated from an environment, excluding “Food”, “Consumer_Product”, and “Biocontrol_Agent” sources; e.g., plants, soil, sewage, ocean, animal feed); (i) “Laboratory” (the strain was used/manipulated by humans in a laboratory experiment and not isolated "naturally" from a source, excluding “Biocontrol_Agent” strains; e.g., genetically modified strains, vaccine strains); (j) “Unknown” (the origin of the strain was unknown).

Genomes assigned to the “Human” and “Animal_Mammal” principal source categories were additionally assigned to a secondary source category: (a) “Clinical” (the strain was isolated from a host clinical sample; e.g., feces, vomit, blood, saliva, etc. from a sick host, outbreak case, a host undergoing treatment for an illness, a deceased host); (b) “Non-Clinical” (the strain was explicitly isolated from the microbiome of a host showing no clinical symptoms/illness; e.g., a fecal sample, skin swab, etc. from a healthy host); (c) “Unspecified” (the strain was isolated from a human or animal host, but it was unclear/ambiguous if the strain was clinical or non-clinical).

***Supplementary Text***

Independent of their isolation source assignments, genomes were further classified based on the following: (a) whether the associated strain was known to be responsible for a human illness/infection (“Yes”, “No”, or “Unknown”); (b) the type of human illness/infection the associated strain was reportedly responsible for (“Anthrax”, “Foodborne_Illness:Emetic”, “Foodborne_Illness:Diarrheal”, “Foodborne_Illness:Unspecified”, “Other”, or “Unknown”); (c) whether the associated strain was known to be responsible for >1 human illness cases (e.g., an outbreak; “Yes”, “No”, or “Unknown”). For (b), strains were categorized as “Anthrax” or “Foodborne_Illness:Emetic” if explicit details were provided linking the strain to an anthrax or emetic foodborne illness case, respectively. Strains were categorized as “Foodborne_Illness:Diarrheal” if diarrhea was reported as the illness type or symptom, in the absence of terms denoting that an illness was emetic-like or anthrax-like (e.g., “emetic”, “vomit”, “anthrax”). “Foodborne_Illness:Unspecified” was used to denote strains that were linked to a foodborne illness case, but it was unclear whether the illness was emetic or diarrheal (e.g., because no additional metadata was provided, or because both emetic and diarrheal symptoms were listed). Strains linked to non-anthrax-like and non-gastrointestinal illnesses/infections were categorized as “Other”. “Unknown” was applied to strains that were either not linked to a human illness case, or to strains with an ambiguous clinical manifestation. If (c) the strain was responsible for an outbreak or cluster of illnesses, a unique identifier was created for the outbreak/cluster, so that all strains isolated in conjunction with the same event could be grouped together. For example, 29 genomes of *B. cereus s.l.* strains isolated in conjunction with an emetic outbreak linked to refried beans in New York state in 2016 were all given the identifier “FBOE:2016USANYSBEANS” (Supplementary Table S1).

Figures showcasing BTyperDB metadata were constructed using the ggplot2 v3.4.4[42] and hchinamap v0.1.0 (https://github.com/czxa/hchinamap) packages in R v4.3.1.[43] Map-based figures with pie charts (i.e., Fig. 3 and Supplementary Fig. S3-S5) were constructed in R v4.1.2 using the following packages: ggrepel v0.9.1 (https://github.com/slowkow/ggrepel), sf v1.0-14,[44,45] scatterpie v0.2.1 (https://cran.r-project.org/web/packages/scatterpie), oz v1.0-22 (https://cran.r-project.org/web/packages/oz), ggplot2 v3.3.6, reshape2 v1.4.4,[46] dplyr v1.0.9 (https://dplyr.tidyverse.org, https://github.com/tidyverse/dplyr), magrittr v2.0.3 (https://magrittr.tidyverse.org, https://github.com/tidyverse/magrittr), sqldf v0.4-11 (https://github.com/ggrothendieck/sqldf, https://groups.google.com/group/sqldf), RSQLite v2.3.3 (https://rsqlite.r-dbi.org, https://github.com/r-dbi/RSQLite), gsubfn v0.7 (https://github.com/ggrothendieck/gsubfn), proto v1.0.0 (https://github.com/hadley/proto), assertthat v0.2.1 (https://cran.r-project.org/web/packages/assertthat), memisc v0.99.31.6 (http://memisc.elff.eu,https://github.com/melff/memisc/), MASS v7.3-55,[47] lattice v0.20-45.[48]

**Database and web portal construction.** To construct the BTyperDB database, two input CSV files were created: (i) one file that describes the column types (e.g., text or number, and whether it should be displayed), (ii) one file that contains all table content. The table content file (ii) was converted into an SQLite3 database for fast querying. The portal back-end provides a RESTful API, implemented using Flask in Python. The front-end was implemented using React, with a map display implemented using the Highcharts library. Nginx was used to direct URLs to the front- and back-end, respectively.

**Species-level phylogeny construction.** To construct a maximum likelihood phylogeny of all 5,976 *B. cereus s.l.* genomes in BTyperDB, Prokka v1.14.6[49] was used to identify and annotate genes in each genome (using default settings and the "Bacteria" kingdom database). GFF files produced by Prokka were supplied as input to Panaroo v1.2.7,[50] which was used to construct a core genome alignment ("-a core") of all 5,976 *B. cereus s.l.* genomes, plus an outgroup genome (the type strain/species representative genome of *B. panaciterrae*; NCBI RefSeq Assembly accession GCF_000430785.1),[51] using the following parameters: (i) "strict" clean mode ("--clean-mode strict"); (ii) MAFFT v7.475[52] as the aligner ("--aligner mafft"); (iii) a core genome sample threshold of 95% ("--core_threshold 0.95"); (iv) a protein family sequence identity threshold of 50% ("-f 0.5"). The resulting core genome alignment was queried using snp-sites v2.5.1[53] using the "-c" and "-C" options to obtain an alignment of core SNPs and the number of constant sites in the alignment, respectively. IQ-TREE v2.2.0.3[54] was used to construct a maximum likelihood phylogeny, using (i) the resulting core SNP alignment as input ("-s"); (ii) an ascertainment bias correction ("-fconst 522883,227567,358674,468428", corresponding to the number of constant sites in the alignment); (iii) the General time reversible (GTR) nucleotide substitution model with invariable sites and FreeRates with 4 categories ("-m GTR+I+R4");[55–57] (iv) one thousand replicates of the ultrafast bootstrap approximation.[58] The resulting unrooted maximum likelihood phylogeny was rooted using the outgroup (using the "root" function in the ape v5.7-1[59] R package with “resolve.root = T”). The rooted *B. cereus s.l.* phylogeny was displayed and annotated in iTOL v6,[60] using metadata from BTyperDB.

All aforementioned steps were repeated to construct species-level phylogenies for GTDB’s (i) *B. tropicus*, (ii) *B. cereus*, and (iii) *B. nitratireducens* species (Supplementary Table S1), using *B. cereus s.l.* str. AH187 (for *B. tropicus* and *B. cereus*) and *B. proteolyticus* str. TD42 (for *B. nitratireducens*) as outgroups (NCBI GenBank Assembly accessions GCA_000021225.1 and GCA_001884065.1, respectively). For each species-level phylogeny, the entire core genome alignment produced by Panaroo (“core_gene_alignment.aln”) was used as input for IQ-TREE (because the core gene alignment contains invariant sites, the “-fconst” option was omitted). Core SNPs produced by Panaroo and snp-sites were additionally supplied to RhierBAPS v1.1.4,[61] which was used to partition each species into clusters using default settings and two clustering levels.

**Pairwise ANI calculations.** ANI values were calculated between all genomes assigned to GTDB’s (i) *B. tropicus*, (ii) *B. cereus*, and (iii) *B. nitratireducens* species using FastANI v1.32 (Supplementary Table S1).[33] Pairwise ANI values were displayed in a heatmap using the pheatmap v1.0.12 R package (https://cran.r-project.org/web/packages/pheatmap/). Dendrograms displayed alongside the heatmaps were constructed as described previously.[32] Briefly, pairwise ANI values were converted to a matrix of symmetric ANI dissimilarities and supplied as input to the “hclust” function in R to perform average linkage hierarchical clustering (for code, see the “ANI.dendrogram” function in the bactaxR v0.2.2[32] R package: https://github.com/lmc297/bactaxR/blob/master/R/anitools.R). For *B. tropicus*, a two-dimensional graph of pairwise ANI values was additionally constructed using the "ANI.graph" function in the bactaxR v0.2.2[32] R package (default settings, except the "ANI_threshold" and "graphout_niter" parameters, which were set to 99 and 1,000,000, respectively).

**Within-lineage variant calling and phylogeny construction.** Snippy v4.6.0 (https://github.com/tseemann/snippy) was used to identify core SNPs within each of the following lineages (selected via holistic consideration of MLST, pairwise ANI values, RhierBAPS clustering, and species-level phylogenomic topology): (i) the ST78 clade, (ii) ST234, and (iii) ST932. For each lineage, a genome or chromosome within the respective lineage was used as a reference genome (i.e., NCBI Nucleotide accession CP009369.1, BTyperDB ID BTDB_2022-0003872.1, and NCBI Nucleotide accession CM000741.1 for the ST78 clade, ST234, and ST932, respectively). For genomes with Illumina sequencing reads available, trimmed paired-end reads were used as input; reads were trimmed using Trimmomatic (using the same parameters and adapters used by Shovill v1.1.0: "ILLUMINACLIP:trimmomatic.fa:1:30:11 LEADING:3 TRAILING:3 MINLEN:30 TOPHRED33"; Supplementary Table S1). For genomes with no linked Illumina sequencing reads, the assembled genome was used as input (Supplementary Table S1). For each lineage, Snippy's default settings were used, and the resulting genome alignment was cleaned using the "snippy-clean_full_aln" command. For each lineage, the resulting cleaned alignment was supplied as input to Gubbins v3.1.3,[62] which was used to identify and remove recombination (using default settings). snp-sites was used to query each resulting recombination-free alignment for core SNPs and constant sites (using the "-c" and "-C" options, respectively). IQ-TREE was used to construct a maximum likelihood phylogeny for each lineage as described above (see section "Species-level phylogeny construction" above), with the following changes: (i) each within-lineage phylogeny was constructed using the optimal nucleotide substitution model selected via ModelFinder[63] (based on their Bayesian information criteria [BIC] values, the TVM+F+I+I+R5, K3Pu+F, and HKY+F models for the ST78 clade, ST234, and ST932, respectively);[64,65] (ii) the following ascertainment bias corrections were applied for each of the ST78 clade, ST234, and ST932: "-fconst 1678491,907290,914922,1677390", "-fconst 1722947,1056733,870014,1832834", and "-fconst 1930650,863811,1152112,1778894", respectively; (iii) phylogenies were midpoint-rooted in iTOL.

**Data availability.** All assembled genomes and metadata generated and curated in this study are publicly available and can be downloaded via BTyperDB (www.btyper.app).

**Code availability.** The BTyperDB source code is open-source and available at https://github.com/henriksson-lab/btyper_website. Additional analyses performed in the study are described in the manuscript text and/or Supplementary Text.

### REFERENCES

***Supplementary Text***


1. Schoch, C. L. *et al.* NCBI Taxonomy: a comprehensive update on curation, resources and tools. *Database* **2020**, (2020).
2. Liu, Y. *et al.* Proposal of nine novel species of the *Bacillus cereus* group. *Int. J. Syst. Evol. Microbiol.* **67**, 2499–2508 (2017).
3. Sternbach, G. The history of anthrax. *J. Emerg. Med.* **24**, 463–467 (2003).
4. Chen, Y. *et al. Bacillus arachidis* sp. nov., Isolated from Peanut Rhizosphere Soil. *Curr. Microbiol.* **79**, 231 (2022).
5. Liu, B. *et al. Bacillus bingmayongensis* sp. nov., isolated from the pit soil of Emperor Qin's Terra-cotta warriors in China. *Antonie Van Leeuwenhoek* **105**, 501–510 (2014).
6. Frankland, G. C. T. & Frankland, P. *Studies on Some New Micro-organisms Obtained from Air*. (1887).
7. Méndez Acevedo, M. *et al.* Novel Effective *Bacillus cereus* Group Species "*Bacillus clarus*" Is Represented by Antibiotic-Producing Strain ATCC 21929 Isolated from Soil. *mSphere* **5**, (2020).
8. Guinebretière, M.-H. *et al. Bacillus cytotoxicus* sp. nov. is a novel thermotolerant species of the *Bacillus cereus* Group occasionally associated with food poisoning. *Int. J. Syst. Evol. Microbiol.* **63**, 31–40 (2013).
9. Liu, X. *et al. Bacillus fungorum* sp. nov., a bacterium isolated from spent mushroom substrate. *Int. J. Syst. Evol. Microbiol.* 70, 1457–1462 (2020).
10. Jung, M. Y. *et al. Bacillus gaemokensis* sp. nov., isolated from foreshore tidal flat sediment from the Yellow Sea. *J. Microbiol.* **48**, 867–871 (2010).
11. Tohya, M. *et al.* Three novel species of the *Bacillus cereus* group isolated from

clinical samples in Japan. *Int. J. Syst. Evol. Microbiol.* **71**, (2021).

12. Jung, M. Y. *et al. Bacillus manliponensis* sp. nov., a new member of the *Bacillus cereus* group isolated from foreshore tidal flat sediment. *J. Microbiol.* **49**, 1027–1032 (2011).
13. Laubach, C. A. Spore-Bearing Bacteria in Dust. *J. Bacteriol.* **1**, 493–505 (1916).
14. Nakamura, L. K. *Bacillus pseudomycoides* sp. nov. *Int. J. Syst. Bacteriol.* **48 Pt 3**, 1031–1035 (1998).
15. Kämpfer, P. *et al. Bacillus rhizoplanae* sp. nov. from maize roots. *Int. J. Syst. Evol. Microbiol.* 72, (2022).
16. Milner, R. J. History of *Bacillus thuringiensis*. *Agric. Ecosyst. Environ.* **49**, 9–13 (1994).
17. Jiménez, G. *et al.* Description of *Bacillus toyonensis* sp. nov., a novel species of the *Bacillus cereus* group, and pairwise genome comparisons of the species of the group by means of ANI calculations. *Syst. Appl. Microbiol.* **36**, 383–391 (2013).
18. Lechner, S. *et al. Bacillus weihenstephanensis* sp. nov. is a new psychrotolerant species of the *Bacillus cereus* group. *Int. J. Syst. Bacteriol.* **48 Pt 4**, 1373–1382 (1998).
19. Miller, R. A. *et al. Bacillus wiedmannii* sp. nov., a psychrotolerant and cytotoxic *Bacillus cereus* group species isolated from dairy foods and dairy environments. *Int. J. Syst. Evol. Microbiol.* **66**, 4744–4753 (2016).
20. Katz, K. *et al.* The Sequence Read Archive: a decade more of explosive growth. *Nucleic Acids Res.* **50**, D387 (2022).
21. Courtot, M., Gupta, D., Liyanage, I., Xu, F. & Burdett, T. BioSamples database:

FAIRer samples metadata to accelerate research data management. *Nucleic Acids Res.* **50**, D1500 (2022).

22. Babraham Bioinformatics. FastQC: A Quality Control tool for High Throughput Sequence Data. https://www.bioinformatics.babraham.ac.uk/projects/fastqc/.
23. Ewels, P., Magnusson, M., Lundin, S. & Käller, M. MultiQC: summarize analysis results for multiple tools and samples in a single report. *Bioinformatics* **32**, 3047–3048 (2016).
24. Bolger, A. M., Lohse, M. & Usadel, B. Trimmomatic: a flexible trimmer for Illumina sequence data. *Bioinformatics* **30**, 2114–2120 (2014).
25. Souvorov, A., Agarwala, R. & Lipman, D. J. SKESA: strategic k-mer extension for scrupulous assemblies. *Genome Biol.* **19**, 153 (2018).
26. Wick, R. R., Judd, L. M., Gorrie, C. L. & Holt, K. E. Unicycler: Resolving bacterial genome assemblies from short and long sequencing reads. *PLoS Comput. Biol.* **13**, e1005595 (2017).
27. Madoroba, E. *et al.* Microbial Communities of Meat and Meat Products: An Exploratory Analysis of the Product Quality and Safety at Selected Enterprises in South Africa. *Microorganisms* **9**, (2021).
28. Gurevich, A., Saveliev, V., Vyahhi, N. & Tesler, G. QUAST: quality assessment tool for genome assemblies. *Bioinformatics* **29**, 1072–1075 (2013).
29. Parks, D. H., Imelfort, M., Skennerton, C. T., Hugenholtz, P. & Tyson, G. W. CheckM: assessing the quality of microbial genomes recovered from isolates, single cells, and metagenomes. *Genome Res.* **25**, 1043–1055 (2015).
30. Carroll, L. M., Cheng, R. A. & Kovac, J. No Assembly Required: Using BTyper3 to

Assess the Congruency of a Proposed Taxonomic Framework for the *Bacillus cereus* Group With Historical Typing Methods. *Front. Microbiol.* **11**, 580691 (2020).

38. Chaumeil, P.-A., Mussig, A. J., Hugenholtz, P. & Parks, D. H. GTDB-Tk v2: memory friendly classification with the genome taxonomy database. *Bioinformatics* **38**, 5315–5316 (2022).
39. Parks, D. H. *et al.* A complete domain-to-species taxonomy for Bacteria and Archaea. *Nat. Biotechnol.* **38**, 1079–1086 (2020).
40. Winter, D. J. rentrez: An R package for the NCBI eUtils API. *R J.* **9**, 520–526 (2017).
41. The R Project for Statistical Computing. (2020) https://www.R-project.org/.
42. Wickham, H. *ggplot2: Elegant Graphics for Data Analysis*. (Springer Science & Business Media, 2009).
43. The R Project for Statistical Computing. (2023) https://www.R-project.org/.
44. Pebesma, E. & Bivand, R. *Spatial Data Science*. (Chapman and Hall/CRC, 2023).
45. Pebesma, E. Simple Features for R: Standardized Support for Spatial Vector Data. *R J.* **10**, 439–446 (2018).
46. Wickham, H. Reshaping Data with the reshape Package. *J. Stat. Softw.* **21**, 1–20 (2007).
47. Venables, W. N. & Ripley, B. D. *Modern Applied Statistics with S*. (Springer Science & Business Media, 2003).
48. Sarkar, D. *Lattice: Multivariate Data Visualization with R*. (Springer Science & Business Media, 2008).
49. Seemann, T. Prokka: rapid prokaryotic genome annotation. *Bioinformatics* **30**, 2068–2069 (2014).
50. Tonkin-Hill, G. *et al.* Producing polished prokaryotic pangenomes with the Panaroo

pipeline. *Genome Biol.* **21**, 180 (2020).

51. Ten, L. N. *et al. Bacillus panaciterrae* sp. nov., isolated from soil of a ginseng field. *Int. J. Syst. Evol. Microbiol.* **56**, 2861–2866 (2006).
52. Katoh, K. & Standley, D. M. MAFFT multiple sequence alignment software version 7: improvements in performance and usability. *Mol. Biol. Evol.* **30**, 772–780 (2013).
53. Page, A. J. *et al.* SNP-sites: rapid efficient extraction of SNPs from multi-FASTA alignments. *Microb Genom* **2**, e000056 (2016).
54. Minh, B. Q. *et al.* IQ-TREE 2: New Models and Efficient Methods for Phylogenetic Inference in the Genomic Era. *Mol. Biol. Evol.* **37**, 1530–1534 (2020).
55. Tavaré, S. Some probabilistic and statistical problems in the analysis of DNA sequences. (1986).
56. Yang, Z. A space-time process model for the evolution of DNA sequences. *Genetics* **139**, 993–1005 (1995).
57. Soubrier, J. *et al.* The influence of rate heterogeneity among sites on the time dependence of molecular rates. *Mol. Biol. Evol.* **29**, 3345–3358 (2012).
58. Hoang, D. T., Chernomor, O., von Haeseler, A., Minh, B. Q. & Vinh, L. S. UFBoot2: Improving the Ultrafast Bootstrap Approximation. *Mol. Biol. Evol.* **35**, 518–522 (2018).
59. Paradis, E. & Schliep, K. ape 5.0: an environment for modern phylogenetics and evolutionary analyses in R. *Bioinformatics* **35**, 526–528 (2019).
60. Letunic, I. & Bork, P. Interactive Tree Of Life (iTOL) v5: an online tool for phylogenetic tree display and annotation. *Nucleic Acids Res.* **49**, W293–W296 (2021).
